# Keeping time with the host: reconstructing the developmental rhythms of malaria parasites

**DOI:** 10.64898/2026.03.25.714289

**Authors:** Zhenying Chen, Lucas A. Nell, Aidan J. O’Donnell, Sarah E. Reece, Megan A. Greischar

## Abstract

Theory predicts that pathogenic organisms benefit from aligning to host circadian rhythms, but observable data provide an incomplete picture of infection rhythms. Models are needed to reconstruct unobservable dynamics and uncover fitness impacts, challenges typified in malaria infections. Malaria parasites often synchronize their development to multiples of 24 hours and reschedule their developmental rhythms following perturbation. Yet it remains uncertain how parasites reschedule—including any cost to parasite multiplication rates—due to the difficulty of detecting parasites in all phases of their development. As parasites mature, the red blood cells they occupy adhere to blood vessel walls (sequestration, an immune evasion tactic) where they cannot be readily sampled. Existing methods cannot determine sequestration timing *in vivo* nor how that timing varies when misaligned to within-host environments. To address these challenges, we fit a mathematical model tracking parasite development and sequestration to high-resolution time series from the rodent malaria parasite *Plasmodium chabaudi*. We show that parasites hasten late development—but not sequestration—to realign with host rhythms. That rescheduling reduces multiplication rates, a result only apparent from model-reconstructed dynamics. Our novel approach recovers the timing of development and sequestration, providing insight into the consequences of circadian rhythms for parasite fitness.

## 1 Introduction

By anticipating and aligning with environmental rhythms, organisms can enhance their survival and reproductive success. Such biological rhythms play critical roles during infection, since within-host environments fluctuation with the time of day [2,3]. The ability of parasites to synchronize with host rhythms should offer significant evolutionary advantages, such as enhancing transmission, optimizing resource exploitation, or evading host immune responses [4, 5].

Among pathogenic organisms, malaria parasites display some of the most conspicuous rhythms [6–9], exhibiting cycles of development and replication within red blood cells (RBCs) that typically correspond to a multiple of 24 hours depending on the *Plasmodium* species [6, 10]. Synchronizing the intra-erythrocytic development cycle (IDC) across the population of parasites within an infection can result in the hallmark periodic fever episodes associated with malaria. The IDC length (period) is not solely host-determined since, for example, malaria parasite species that infect humans exhibit IDCs of either 24, 48, or 72 hours [11]. IDC timing and synchronization is predicted to impact parasite ecology within the host, including transmission success [12], a prediction supported by experiments [5]. Rodent malaria *P. chabaudi* infections suggest both within-host multiplication and onward transmission decline when parasite development is misaligned with host circadian rhythms (parasite ‘jet lag’, [13]). IDC scheduling also impacts resilience to antimalarial drug treatment [14] with delayed parasite development implicated in the drug resistance phenotype of human malaria (*P. falciparum*) parasites to the current front-line drug, artemisinin ([15], reviewed in [9]).

Understanding how and why malaria parasites align their developmental schedules to host rhythms requires overcoming a major methodological challenge: capacity to accurately quantify within-host abundance depends on parasite developmental schedules (developmental sampling bias, [16]). As parasites mature, the infected RBCs (iRBCs) they inhabit tend to stop circulating and adhere to the walls of blood vessels and capillaries (“sequester”), becoming undetectable in blood samples (e.g., [10]). This aspect of *Plasmodium* biology is shared with other Apicomplexans including *Babesia* [17] and *Trypanosoma* [18], but existing methods cannot reveal the timing or extent to which parasites disappear from view *in vivo*. That represents a key gap in understanding since sequestration impacts disease severity, both directly when sequestered iRBCs hinder blood flow to vital organs, and indirectly by obscuring health risks when parasite biomass is dangerously high but largely hidden from view (reviewed in [19,20]). For sequestration to occur, parasites must express specific proteins that enable adhesion to host endothelial cells. The developmental timing of protein expression can be quantified in artificial culture where such methods exist (e.g., for *P. falciparum*, [21]). However, insight is limited by the lack of artificial culture methods for most *Plasmodium* species [22]. Further, sequestration is only possible when the host expresses corresponding receptors to allow parasite attachment (reviewed in [23]). Rodent malaria infections showed suppressed parasite population growth when sequestration was hindered by genetic modifications to either parasites [24] or hosts [25], but links between sequestration and within-host dynamics remain uncertain in the vast majority of host-parasite combinations. By necessity, efforts to reconstruct total within-host abundance (circulating and sequestered) rely on assumptions about synchrony and IDC timing derived from *in vitro* data (where available) [26] with unknown relevance to *in vivo* conditions (reviewed in [27]). Therefore, methods are needed that can recover the timing of sequestration *in vivo* to address questions about parasite biology, including the adaptive significance of IDC rhythms.

Parasite fitness consequences can be revealed by perturbing IDC rhythms relative to host rhythms, feasible through experimental *P. chabaudi* infections of mice. A recent study [1] investigated parasite dynamics after synchronized parasites from donor mice were inoculated into wild type (WT) mice with a circadian rhythm matched or mismatched to donor mice, or genetically-modified clock-disrupted mice fed on a mismatched rhythm or continuously to damp IDC rhythms. Parasites shortened their IDC to reschedule when misaligned to host rhythms, but with no detectable reduction in circulating iRBC abundance. We develop a model-fitting approach to reconstruct within-host dynamics despite sequestration and address two major unknowns using the data from [1]: 1) how parasites reschedule their development—especially sequestration timing—to align with host rhythms, and 2) how rescheduling modifies parasite multiplication rates (PMRs), a key component of parasite fitness. We find that *P. chabaudi* sequesters remarkably late in its development (≈19 hours into the 24 hour IDC), with no evidence that misalignment to host rhythms alters sequestration timing. This timing is strikingly similar to patterns derived *in vitro* for the human *P. falciparum* parasites [21], despite the fact that *P. falciparum* has a 48-hour IDC compared to the 24-hour IDC of *P. chabaudi*. Accounting for sequestration reveals that parasites suffer a fitness cost—significantly reduced PMRs—when misaligned to host rhythms, a result not evident from observed data alone. The approach provides, for the first time, a method to reconstruct sequestration patterns and within-host dynamics *in vivo* and understand how parasite developmental schedules interact with immunity, anti-malarial drug sensitivity and disease severity.

## 2 Methods

We adapted a model framework previously used for human *P. falciparum* infections [16,28,29] to experimental *P. chabaudi* infections of mice. The model uses ordinary differential equations to track parasite development within iRBCs through two parallel tracks, circulating and sequestered, enabling simulation of circulating and total iRBC abundance over the course of infection. We extend that framework to simulate the percentage of circulating iRBCs containing parasites in the early stages of development (known as the ‘ring stage’ due to the ring-like appearance of parasites in that phase of development). The percentage of circulating iRBCs in the ring stage represents a commonly used but unacceptably biased metric of synchrony, since asynchronous populations can exhibit a large percentage of ring stage parasites due to rapid expansion of the iRBC population [9,30]. However, synchrony can be quantified by total iRBC abundance in the absence of sequestration [30]. To examine dynamics with sequestration, we use the model to be fitted—which incorporates sequestration—to identify the impact of modulating different parameter values on simulated dynamics of circulating iRBCs and ring percentages (i.e., the percentage of circulating iRBCs in the ring stage of development). That analysis shows that ring percentages can provide useful information when coupled with circulating iRBC time series. Three parameters emerged as important to both circulating iRBC and ring percentage dynamics, parasite developmental age at 50% sequestration, IDC duration, and ring stage duration. Focusing on those key parameters, we fit the model to high-resolution time series data from experimental *Plasmodium chabaudi* infections of mice [1] in standard conditions or in which parasites were misaligned to host rhythms. We fit to each treatment group, using forward model selection to determine when fitting—rather than fixing—additional parameters gave a significantly better fit to the data. We used parametric bootstrapping to construct confidence intervals for each fitted parameter from 100 synthetic datasets that preserve the mean and variance at each sampling time point.

### 2.1 Experimental rodent infection data

Full details of the experimental methods can be found in the original study [1]. Briefly, wild-type (WT) mice (C57BL/6J) were acclimatized to a standard 12 hours light–12 hours dark (LD) photoperiod with food available during the dark phase and then infected with *P. chabaudi* DK strain to use as donors for subsequent infections. The initial inoculum was highly synchronized by sampling from donor mice when nearly all circulating iRBCs contained ring stage parasites. Pooled samples from three donor mice were used to infect recipient mice that were either WT or canonical transcription-translation feedback loop (TTFL) clock-disrupted, behaviorally arrhythmic (*Per1/2*-null). Since host feeding rhythms have shown to have a significant influence in driving parasite developmental schedules [7], mice either experienced time restricted feeding (TRF, food available for 12 hours per day) or food was provided at all times. Together, there were a total of four treatment groups (Fig. 1a): (1) WT mice with *ad libitum* food which they primarily eat at night, following same photoschedule as the donor mice (WT matched group), (2) WT mice with *ad libitum* food acclimated to a photoschedule 12 hours offset from the donor mice (WT mismatched group), (3) *Per1/2*-null mice kept in constant darkness to maintain behavioral arrhythmia, with time restricted feeding (TRF) so that food was available for 12 hours each day corresponding to feeding rhythms in the WT mismatched group (*Per1/2*-null TRF), and (4) *Per1/2*-null mice kept in constant darkness to maintain behavioral arrhythmia with continuous food availability to remove feeding rhythms (*Per1/2*-null all-day fed).

**Figure 1:**
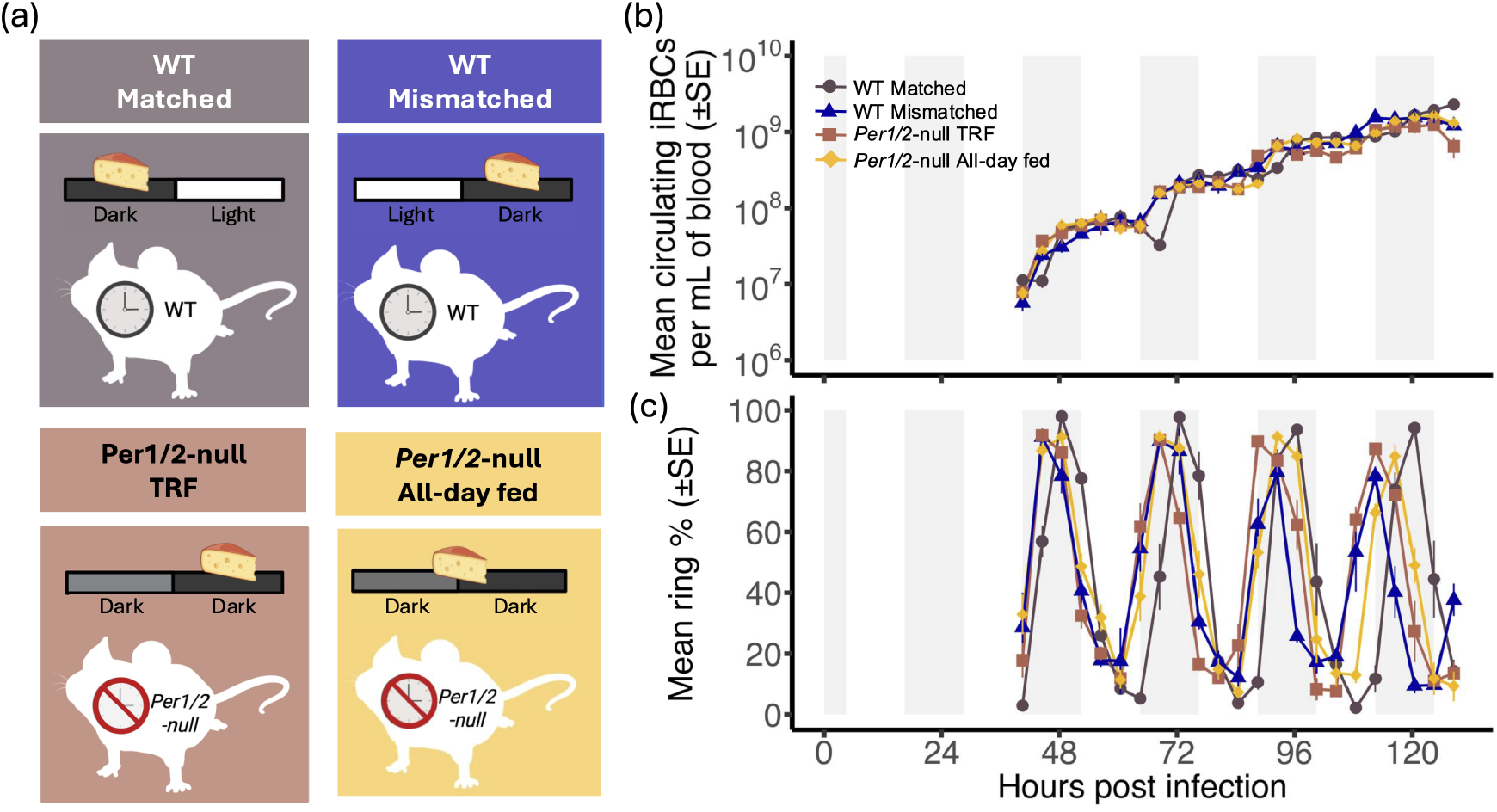
*P. chabaudi* exhibits altered circulating iRBCs and ring stage percentage dynamics in perturbed within-host environments. (*a*) The four treatment groups in [1] are shown with backgrounds colored to match the corresponding dynamics shown at right (see main text for details of treatment groups). Mice were either wild-type (“WT”) or genetically modified “*Per1/2*-null” as shown by the text in each mouse outline. Daily 12 hour periods of light or dark are indicated with white or dark gray rectangles above each mouse, with a slice of cheese to indicate the 12 hour window when mice fed (WT Matched, WT Mismatched and *Per1/2*-null TRF) or spanning the two 12 hour periods to denote continuous food availability (*Per1/2*-null all-day fed). Here, “matched” and “mismatched” indicate recipient mice were respectively either on the same circadian schedule as the donor mice or a 12 hour phase shift from donor to recipient mice.(*b*) The abundance of mean circulating iRBCs per mL increases in a log-linear fashion during the initial expansion phase for WT matched (circles), WT mismatched (triangles), *Per1/2*-null TRF (squares), and *Per1/2*-null all-day fed (diamonds) groups. Grey vertical boxes indicate periods when WT donors were in the dark phase of the photoperiod. (*c*) The percent of circulating iRBCs in the ring stage of development fluctuates periodically for each treatment group (colors and symbols as in B), with perturbed treatments exhibiting faster oscillations than the WT matched group.

Following infection of recipient mice, samples were taken every 4 hours starting at 40.5 hours post infection. For each sample, the abundance of RBCs per mL of blood was measured by flow cytometry and a blood smear was prepared to determine the proportion of RBCs infected via microscopy. Approximately 100 iRBCs were used to estimate the percentage of ring stage iRBCs per time point. Due to practical and ethical constraints on how frequently blood can be sampled—e.g., sampling too frequently could perturb parasite rhythms—mice were separated into four cohorts and each cohort was sampled for a different 28 hour-window. These data were concatenated into continuous time series that we used to fit the model (Fig. 1b,c).

The time series collected by [1] includes an initial log-linear expansion in iRBC numbers sampled during days 2-6 until peak iRBC abundance, followed by declining iRBC abundance during the post-peak window sampled on days 7-10. We focus on the initial log-linear expansion phase of infection, since the model framework we adapt (details below) assumes a constant PMR. To determine when the initial expansion phase ends, we smooth circulating iRBC abundance for all 4 treatments and define the cutoff time as the point at which the derivative of smoothed abundance reaches zero, indicating that the infection has reached peak parasite abundance (see Supplemental methods S1.1, Fig. S1). Based on this criterion, the cutoff time for the acute phase across all treatment groups is 128.5 hours post infection.

### 2.2 Model-fitting approach

We used an ordinary differential equation (ODE) model modified from [28] that tracks parasite development through *n* age classes in two parallel tracks (circulating and sequestered, see Fig. 2 and Table 1 for parameter definitions). We denote the circulating iRBCs in developmental age class *i* by *C*_*i*_, where *C*_1_ represents newly invaded iRBCs and *C*_*n*_ represents fully matured circulating iRBCs that are ready to burst and initiate a new infection cycle. Similarly, *S*_*i*_ indicates sequestered iRBCs in age class *i*, where *S*_1_ represents newly invaded sequestered iRBCs and *S*_*n*_ represents fully matured sequestered parasites:

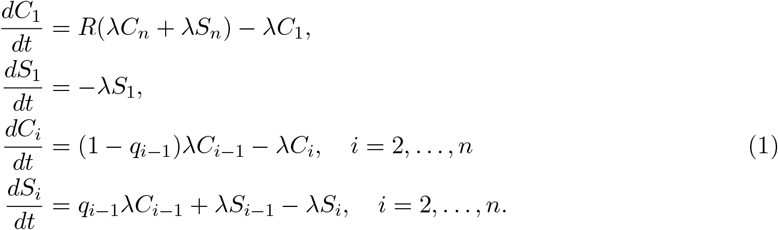

**Figure 2:**
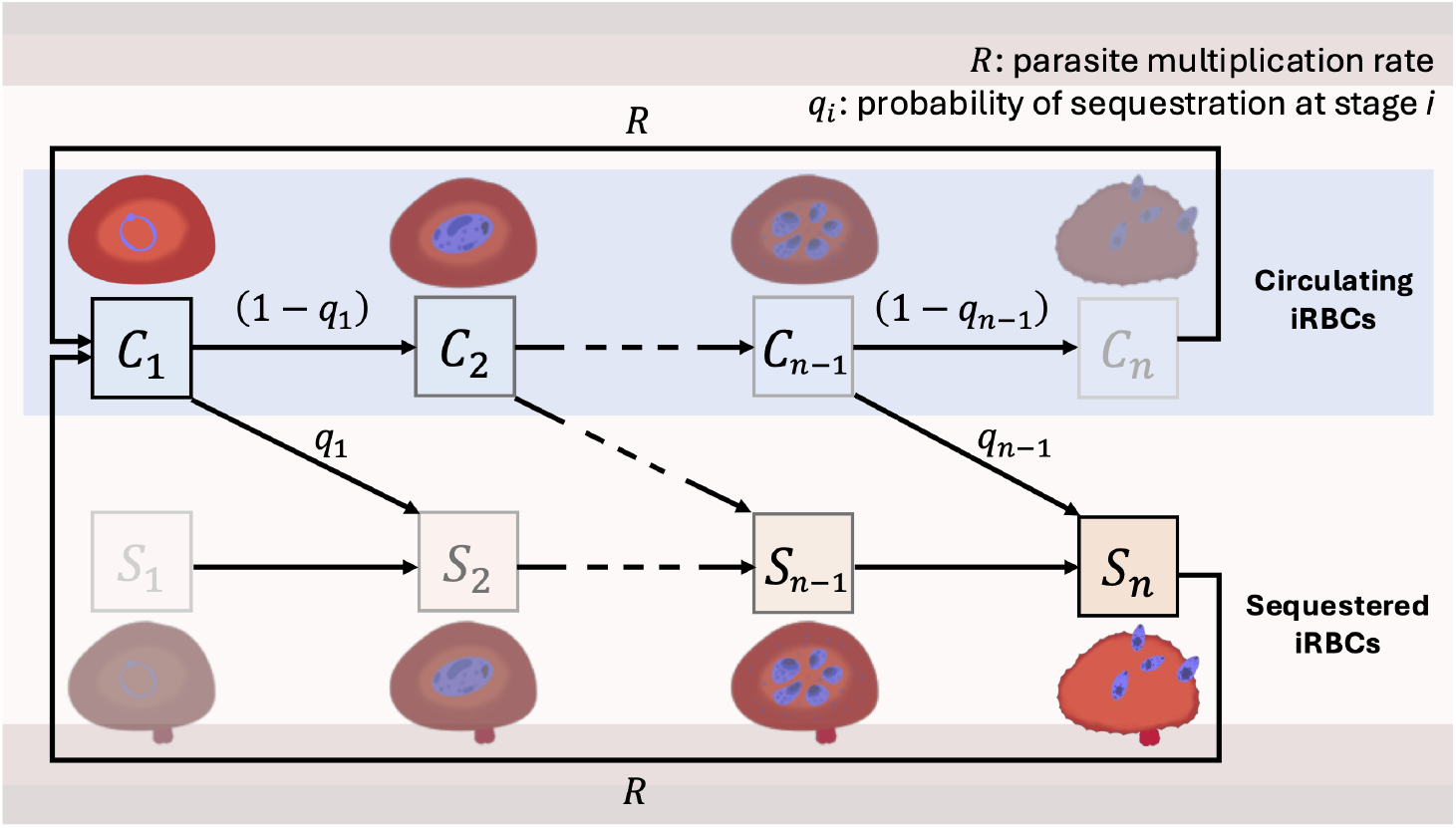
The model tracks parasite development within iRBCs with transitions from circulating to sequestered. In the blood, parasites invade and mature within host RBCs. Parasites begin development within circulating iRBCs (blue box) *C*_*i*_ and, as they mature, become increasingly likely to transition to sequestered iRBCs *S*_*i*_. Transparent age compartments indicates that the parasites are unlikely to be in that stage (e.g., iRBCs sequestering at the beginning of development or iRBC continuing to circulate late in development). Circulating iRBCs *C*_*i*_ and sequestered iRBCs *S*_*i*_ mature at a rate *ω*. Parasites progress through *n* age classes, with the fraction of iRBCs sequestering at stage *i* denoted by *q*_*i*_. Upon completing development, each iRBC bursts to generate *R* new circulating iRBCs. For clarity, only parameters associated with parasite proliferation and sequestration are shown (see Methods for full model).

**Table 1:** Parameters used in the blood stage model of parasite development.

| Parameters | Description | Value | Unit | Ref |
| --- | --- | --- | --- | --- |
| $R$ | Parasite multiplication rate | fitted | new iRBCs/bursting iRBC | - |
| $q_i$ | fraction of iRBCs sequestering at stage $i$ | $1 - \frac{g_i}{g_{i-1}}$ | - | [21, 31] |
| $c$ | IDC duration | fitted(20 – 28) | hr | - |
| $n$ | Number of developmental age compartments | fitted | - | - |
| $I_0$ | Inoculum size of iRBCs | fitted | /mL of blood | - |
| $o$ | Offset of initial iRBCs distribution, specifies initial median parasite developmental age (hour) | fitted | - | - |
| $s$ | Shape of the initial distribution of iRBCs | fitted | - | - |
| $r_L$ | Ring stage duration | fitted | hr | - |
| $p_1$ | Minimal sequestration protein expression level | 11.3869/467.6209 | - | [?, 31] |
| $p_2$ | Maximum sequestration protein expression level | 1 | - | [?, 31] |
| $p_3$ | Inflection point of the probability of sequestration curve | fitted | hr | - |
| $p_4$ | Steepness of the probability of sequestration curve at inflection point | $0.2242 \cdot 2$ | - | [?, 31] |

Both circulating and sequestered iRBCs mature at a rate *ω* = *n/c*, where *c* represents the IDC duration in hours. As parasites mature within circulating iRBCs *C*_*i*_, a fraction 1 *− q*_*i*_ remains in circulation, while a fraction *q*_*i*_ sequester to host capillary walls. Any attrition of iRBCs is incorporated into the PMR (*R*), which represents the average number of new iRBCs produced per bursting iRBC. Thus, reductions in *R* could indicate any combination of decreased survival of iRBCs, fewer RBC-invasive daughter cells (merozoites) produced per iRBC, or reduced invasion success. During the pre-peak phase of infection, iRBC population growth is approximately exponential (i.e., log-linear). Therefore, we assumed the PMR *R* was a nonnegative constant, an assumption we later revisit (see “Identifying the signal of variation in model parameters” below).

We model *q*_*i*_, the fraction of cells that mature from a circulating developmental age class *I* into a sequestered age class *i* +1 as

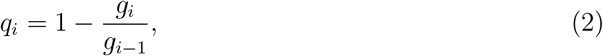

where *g*_*i*_ denotes the scaled rate at which iRBCs in age class *i* transition to the sequestered developmental track. We retain the same formulation as [28], which enables utilization of *in vitro* data on the developmental timing of production of proteins required for sequestration. As parasites mature within RBCs, they express proteins to sequester to walls of blood vessels and become undetectable in blood samples (e.g., [10]), so we assume that the scaled rate of sequestration (*g*) increases with *t*, the developmental age of parasites within iRBCs following a sigmoidal function defined by [28]:

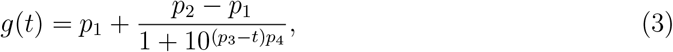

where *p*_1_ and *p*_2_ are the lower and upper bounds of the curve, *p*_3_ is the inflection point corresponding to the parasite age at which the probability of sequestration is 50%, and *p*_4_ is the slope at the inflection point.

The simulated abundance of circulating iRBCs is defined as

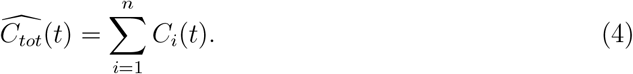

To calculate the percentage of circulating iRBCs in the ring stage, we extend the models used in [16, 28, 29] by introducing an additional parameter representing the duration of ring stage development, *r*_*L*_. Simulated ring percentages 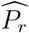 are calculated as:

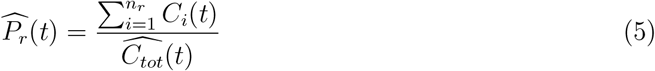

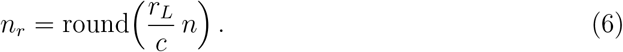

The decay of synchrony is determined by the starting synchronization (i.e., the age structure of the initial inoculum of iRBCs) and the variability in IDC duration (a function of the number of age compartments *n*). We divide the total initial abundance of iRBCs (*I*_0_) among *n* developmental age classes according to a symmetric Beta distribution as in [16, 29]. The shape parameter *s* determines the initial level of synchronization, ranging from asynchrony (*s* = 1) to high levels of synchrony (i.e., large *s* values yield extremely narrow bell curves in the age distribution). By default, the median of a symmetric Beta distribution falls in the center (0.5 × the cycle length *c*), so we follow [16, 29, 30] and introduce an offset parameter *o* to allow the initial median parasite developmental age to deviate from the default.

### 2.3 Identifying the signal of variation in model parameters

We simulated the dynamics of circulating iRBCs and ring percentages by increasing or decreasing one parameter at a time while keeping all others fixed. We varied parameters controlling parasite development and initial conditions (*c, I*_0_, *o, s*), four parameters defining the probability of sequestration curve (*p*_1_, *p*_2_, *p*_3_, and *p*_4_), and ring duration *r*_*L*_. Unless otherwise stated, baseline parameter values were *n* = 100 (number of developmental age compartments), *R* = 3 (parasite multiplication rate), *I*_0_ = 10^6^ (initial abundance of iRBCs), *o* = 0 (offset of the initial iRBC age distribution; *o* = 0 equates to an initial median parasite developmental age of 12 hours), *s* = 30 (shape of the initial iRBC age distribution; *s* = 30 translates into an age distribution with a width of 8 hours (i.e., the difference between the 99.5% and 0.5% quantiles of the age distribution, [30]), *c* = 24 hours (IDC duration), and *r*_*L*_ = 6 hours (ring stage duration). Parameters governing the probability of sequestration were set to *p*_1_ = 0.024, *p*_2_ = 1 (minimal and maximal sequestration level reported for *P. falciparum in vitro*, [21]), *p*_3_ = 12 (inflection point), and *p*_4_ = 0.4484 (steepness at the inflection point). Parameters *p*_1_, *p*_2_, and *p*_4_ were taken from the fitted probability of sequestration curve for *P. falciparum*, while *p*_3_ was set to 12 (halfway through the IDC).

Since iRBC population growth slows during the pre-peak phase of infection (Fig. S1), we also investigated the impact of assuming constant PMR by redefining *R* as a decreasing linear function of hours post-infection to allow parasite population growth to decline towards the peak of infection. In addition to examining the impact of constant versus decreasing PMRs on observable dynamics, this comparison provided information regarding the consequences of a key model assumption—constant PMR—on estimated initial conditions for which data are unavailable.

### 2.4 Fitting & model selection

We defined the best fit parameter values as those yielding the lowest sum squared error (SSE):

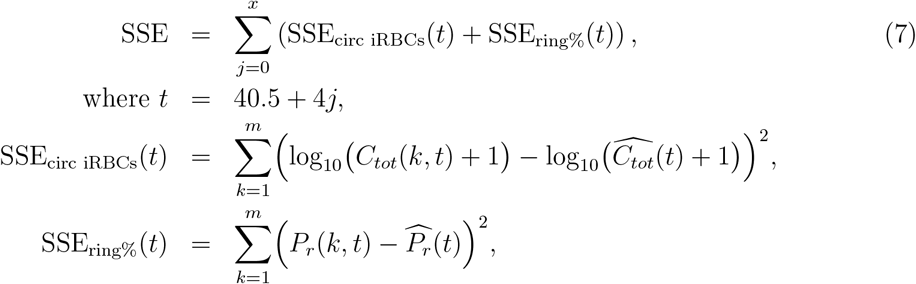

where the squared errors are summed across *x* = 22 samples for *m* replicate mice (*m* = 4 or 8, depending on the time point, see Methods for details). The observed circulating iRBC abundance at hour post-infection *t* for mouse *k* is denoted *C*_*tot*_(*k, t*), with paired data on the percentage rings indicated by *P*_*r*_(*k, t*). The model predicted values 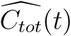 and 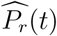 are defined in Eq. 4 and 6. To reduce computational time while increasing the odds of locating a global best fit set of parameter values, we used a winnow optimization algorithm (details in Supplemental Methods S1.2), in which we set bounds on the biologically plausible range for each fitted parameter value (see Table S1).

We employed forward model selection, beginning with a basic five-parameter model fitting the number of developmental age compartments *n*, parasite multiplication rate *R*, median age of the initial iRBC distribution *o*, shape parameter of the initial iRBC distribution *s*, and parasite inoculum size *I*_0_. We retained the same lower and upper bounds on sequestration probability (*p*_1_ and *p*_2_) as those fitted for *P. falciparum*, corresponding to near-zero sequestration in newly invaded iRBCs and complete sequestration in fully matured iRBCs (Table 1). To fit the WT matched data, we set the inflection point for sequestration *p*_3_ = 9.29 hours initially, half the *p*_3_ = 18.58 hour value reported for *P. falciparum in vitro* [21, 31] *to reflect that the IDC of P. chabaudi* is half as long. The ring stage duration *r*_*L*_ was initially assumed to be 12 hours based on previous literature suggesting that parasite populations spend approximately half the IDC in the ring stage [32, 33].

We then sequentially increased model complexity by fitting one additional parameter at a time focusing on parameters that alter circulating iRBC abundance and ring percentages (see Methods 2.3), using F-tests to assess whether the each additional parameter significantly improved model fit (SSE). The model selection process for the treatment groups (WT mismatched, *Per1/2*-null TRF, and *Per1/2*-null all-day fed) followed a similar process, save that we set the default parameter values to the best fit model for the WT matched data (the null hypothesis). Thus, the inflection point of probability of sequestration curve *p*_3_, the slope of inflection point *p*_4_, and the ring stage duration *r*_*L*_ were all set to the corresponding fitted parameter values from WT matched group.

Circulating iRBC abundance and ring percentages were reported from day 2 onwards due to the infeasibility of sampling small numbers of iRBCs initially present in infections. We simulated the model from 0 hours post-infection, enabling extrapolation of dynamics prior to data collection. Though no data were collected during this early period, two pieces of information are available about initial conditions: (1) that 10^6^ iRBCs were injected into recipient mice, with some minor, unavoidable sampling error; and (2) that these iRBCs would have been circulating and in the ring stage, since the inoculum was collected when the circulating iRBC populations in donor mice were at nearly 100% rings. We compared alignment between best fit model dynamics and the expected initial conditions to gain insight into the processes at play during the earliest days of infection.

### 2.5 Parametric bootstrapping for generating confidence intervals

To obtain confidence intervals for fitted parameters, we used parametric bootstrapping following [34] and generated 100 synthetic datasets from the circulating iRBCs and ring stage percentage time series that preserve the mean and variance of the observed data at each time point. Because iRBC abundance measurements were obtained from different mice at each time point, the mean-variance relationship cannot be directly observed. Instead, we fit the relationship between the mean (*µ*) and variance (*σ*^2^) of WT matched circulating iRBC abundance on a log scale at each time point using:

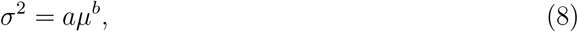

The fitted exponent was close to 2 (*b* = 1.884), consistent with a Gamma-distributed error and very close to values previously estimated by [34] for quantitative PCR counts of *P. chabaudi*. Following [34], we simplified by fixing *b* = 2, which still provided a good fit to the empirical mean-variance relationship (Fig. S2, see Supplemental methods S1.3 for more details). Refitting under this assumption yielded an estimated dispersion parameter of *a* = 0.106.

For each synthetic iRBC abundance dataset, we also generated corresponding synthetic ring percentage data (Fig. S3). In the original experiment, ring percentages were estimated by examining blood smears and counting approximately 100 iRBCs per sample. Accordingly, for each time point *t* post infection, synthetic ring percentages were generated using a binomial distribution with sample size 100 and the probability set to the mean ring percentage in the original data at that time point. We used each of these replicate synthetic datasets to refit the ODE model and estimate the confidence intervals for fitted parameters.

## 3 Results

### 3.1 Model dynamics depend on the PMR and the timing of sequestration & development

During the first three days of simulated infections, four parameters played a dominant role in shaping circulating iRBC abundance and the mean ring percentage: PMR *R*, the inflection point of the percentage sequestering with developmental age *p*_3_, the parasite IDC duration *c*, and the ring stage duration *r*_*L*_ (Fig. 3). PMR *R* primarily influences the dynamics of circulating iRBCs, with a comparatively smaller effect on ring percentages (Fig. 3*a*). Higher PMR *R* values lead to more rapid increases in circulating iRBCs and shifts the iRBC age distribution toward younger ages, resulting in a greater proportion of detectable ring-stage iRBCs and earlier peaks in ring percentages.

**Figure 3:**
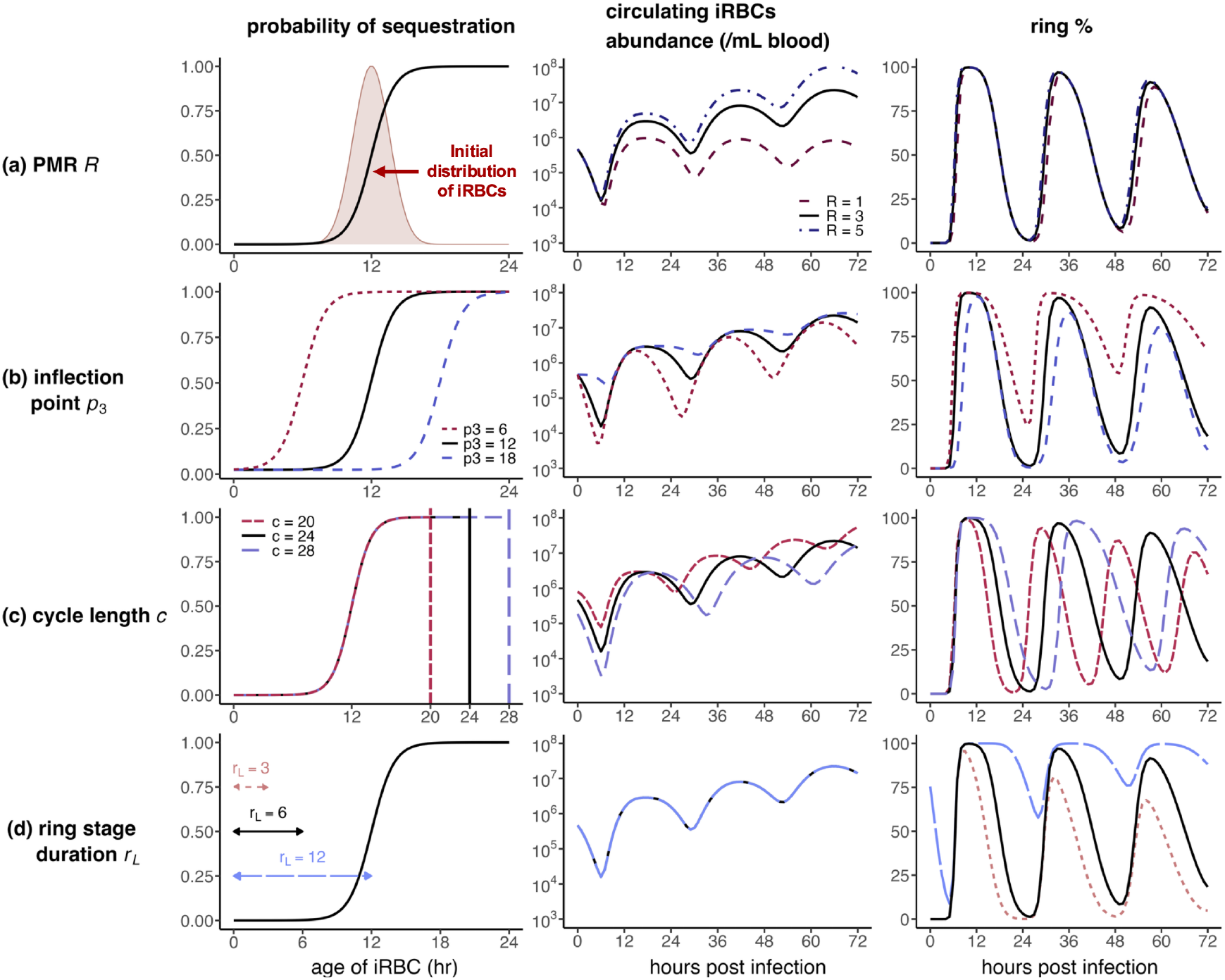
Variation in PMRs and the timing of development and sequestration manifests as changes in observable time series. Left panels show how the probability of sequestration changes with parasite developmental age, while middle panels shown corresponding differences in circulating iRBCs, and right panels show associated ring percentage dynamics. (*a*) Relative to the default scenario (*R* = 3, black solid line), a lower PMR (*R* = 1, red dashed line) reduces the magnitude of circulating iRBC abundance and slightly delays peaks in ring percentages. Conversely, a higher PMR (*R* = 5, blue dot-dashed line) increases the magnitude of circulating iRBC abundance and hastens peaks in ring percentages. (*b*) Relative to the default sequestration curve (*p*_3_ = 12, black solid line), earlier sequestration (*p*_3_ = 6, red short dashed line) increases oscillations in circulating iRBCs while dampening ring stage oscillations, whereas later sequestration (*p*_3_ = 18, blue long dashed line) produces the opposite effect. (*c*) Cycle length *c* shifts the timing and magnitude of circulating iRBC oscillations. Shorter IDC duration (*c* = 20, red short dashed lines) advances and reduces oscillations in circulating iRBC abundance, whereas longer IDC duration (*c* = 28, blue long dashed lines) delays and amplifies those oscillations. Vertical lines in left panel indicate cycle length cutoff times for the corresponding sequestration curves. (*d*) Ring stage duration *r*_*L*_ has no impact on circulating iRBC abundance. Short ring stage duration (*r*_*L*_ = 3, red dotted lines) reduces peak ring percentages, while longer ring stage duration (*r*_*L*_ = 12, blue long dashed lines) maintains high ring percentages. Arrows in left panel indicate ring stage duration in each case. Baseline parameters are *p*_1_ = 0.024, *p*_2_ = 1, *p*_3_ = 12, *p*_4_ = 0.4484, *n* = 100, *R* = 3, *I*_0_ = 10^6^, *o* = 0, *s* = 30, *c* = 24, and *r*_*L*_ = 6.

Smaller values of inflection point *p*_3_ (the parasite developmental age at which sequestration probability is 50 percent) correspond to earlier sequestration during the IDC, resulting in a greater overall percentage of iRBCs being sequestered rather than circulating. As a consequence, circulating iRBC dynamics exhibit larger oscillations within each IDC, reflecting earlier transitions between circulating and sequestered parasite populations. When the inflection point *p*_3_ is early, only the early developmental stage (ring stage) will be circulating and the later stages will almost always be sequestering, so the oscillations of ring percentage never reach zero percent. As synchrony in parasite bursting gradually decays over successive IDCs, the age distribution of the iRBCs becomes wider so that the percentage of circulating ring stage iRBCs remains consistently high. In contrast, larger values of the inflection point *p*_3_ delay sequestration to later stages of the IDC, leading to a higher fraction of mature parasites remaining in circulation, smoother circulating iRBCs dynamics, and larger-amplitude oscillations in ring percentages (Fig. 3b).

We assumed that changes in cycle length (*c*) do not alter the shape of the sequestration probability curve, but instead truncate or extend the developmental timeline. For a given inflection point *p*_3_, increasing the cycle length allows parasites to remain sequestered for a greater portion of their development, leading to larger-amplitude oscillations and delayed peaks in circulating iRBC dynamics. In contrast, shortening the cycle reduces the proportion of development available for sequestration, producing earlier transitions (Fig. 3c)). For ring percentages, changes in cycle length extend or contract the periodicity of oscillations without altering the amplitude.

While ring stage duration *r*_*L*_ has no effect on circulating iRBC abundance, it strongly influences the magnitude of oscillations in ring stage percentage (Fig. 3d). Longer ring stage durations increase the likelihood that circulating iRBCs are observed in the ring stage, preventing the ring percentage from reaching low troughs. Conversely, when ring stage duration *r*_*L*_ is short, parasites rapidly transition out of the ring stage, resulting in lower peak ring percentages and preventing the system from reaching near-complete ring dominance during later cycles.

Of the three parameters governing initial conditions, the level of synchronization (*s*) and inoculum size (*o*) mainly alter fluctuations in circulating iRBCs with little impact on ring percentages (Fig. S4 a, b). We found that increasing *s* mainly affects early infection dynamics prior to the first observed time point, with negligible effects over the data-supported time window (Fig. S5). Since larger *s* values greatly extend the time required to simulate the model, we choose an upper bound of *s* = 300 in subsequent model fitting. The initial median age (*o*) changes the phase of fluctuations in circulating iRBCs and ring percentages, without modifying the trajectories themselves (Fig. S4c). Thus, the three parameters determining initial conditions should be estimable from the high resolution data from [1]. The three other parameters governing sequestration—the upper bound *p*_2_, the lower bound *p*_1_ and the slope at the inflection point *p*_4_—have very little impact on the dynamics of circulating iRBCs or percentage rings (Fig. S6), so we retain our baseline assumed values for these parameters (see Methods).

To revisit our assumption of constant population growth, we also considered a declining PMR constrained to have the same average value as the constant PMR. Both assumptions produced nearly identical predictions for circulating iRBC dynamics and ring-stage proportions for the time period for which data are available (Fig. S7), so we retain the assumption of constant PMR to simplify model fitting and enable direct comparison across treatment groups. However, decaying versus constant PMRs yield distinct dynamics in the phase prior to data collection, suggesting that assuming constant PMRs will lead us to overestimate the initial inoculum size and—to a lesser degree—the initial median age of iRBCs.

### 3.2 Parasites sequester late in development

We began fitting a 5-parameter model to the control (WT matched) data, resulting in estimates for the number of age compartments *n*, the PMR *R*, the initial median age *o*, inoculum size *I*_0_ and synchronization *s*. To determine whether more complicated models were justified by the data, we fitted three 6-parameter models consisting of the 5 baseline parameters initially fitted, with an additional fitted parameter for either cycle length (*c*), the inflection point of sequestration (*p*_3_) or ring duration (*r*_*L*_). Fitting cycle length returned an estimate of almost exactly 24 hours and gave no significant improvement of fit (Table S2), so we retained our baseline assumption of a 24 hour IDC for the control group. Both that baseline assumption and the fitted cycle length align well with previous estimates using a range of approaches on these data [1].

Assuming a 24 hour IDC duration, fitting either the inflection point *p*_3_ or the ring duration *r*_*L*_ gave a significantly better fit to data but including *p*_3_ or *p*_*L*_ caused these parameters to approach their upper or lower plausible bound, respectively (Table S2). Fixing ring duration and fitting the inflection point yielded oscillations in ring percentage noticeably wider than those observed in the data (Fig. S8a, b), while fixing the inflection point and fitting ring duration could not recover the amplitude of oscillations observed in the ring percentage data (Fig. S8c, d). We therefore fit both the ring duration and inflection point (in addition to the baseline 5 parameters) in a 7-parameter model. The best fit 7-parameter model recapitulated the observed ring percentage dynamics (Fig. 4a) and yielded a significantly improved fit compared to either 6-parameter model (Table S2). Hence the best model fit suggests that individual iRBCs take approximately 6 hours—instead of our initially assumed 12 hours—to complete ring stage development and that 50% of iRBCs have sequestered by about 19 hours into their development, instead of our initial assumption of 9 hours (Fig. 4b). The fitted inflection point for sequestration (*p*_3_ = 19.09 hours) is similar to the 18.58 hours estimated for *P. falciparum in vitro* [28] despite the two-fold differences in IDC duration.

**Figure 4:**
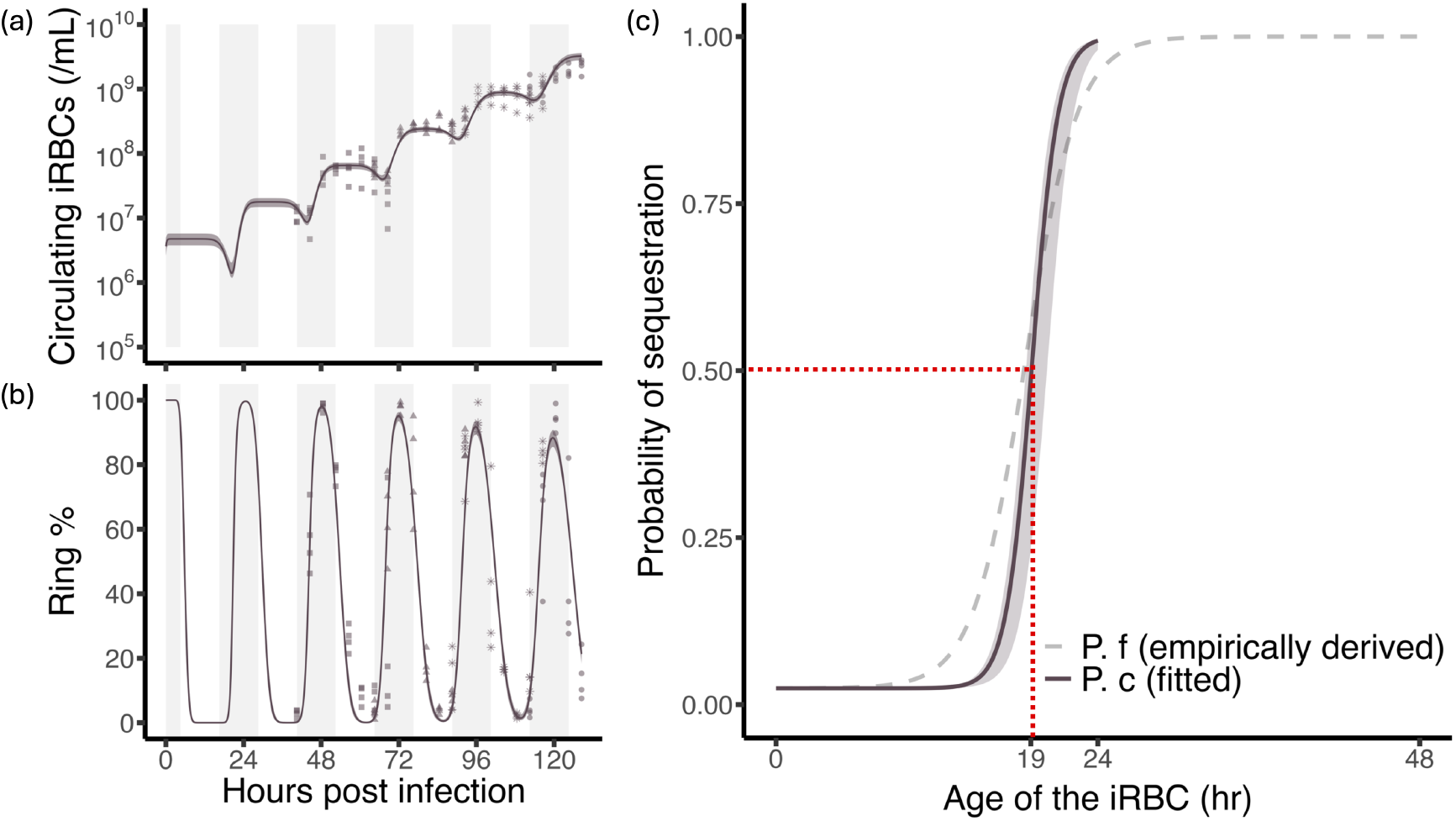
Model-fitted trajectories capture observed within-host dynamics and suggest that *P. chabaudi* sequesters late in the IDC with similar timing to *P. falciparum*. (*a*) Best-fitting model trajectories for circulating iRBC abundance (upper panel) and (*b*) ring percentage are shown for the WT matched (solid lines). Bands with around solid lines show 95% confidence intervals for the dynamics fitted to datasets simulated through parametric bootstrapping. Empirical data are shown in the background, with point shapes indicating different mouse cohorts sampled at each time point (details in [1]). Grey vertical boxes indicate periods corresponding to the dark phase of the photoperiod in WT matched hosts. (*c*) *P. chabaudi* and *P. falciparum* exhibit remarkably similar sequestration timing despite large differences in life-cycle length. The empirically-derived sequestration probability curve for *P. falciparum* (grey dashed line [**?**, 31]) is nearly identical to the model-reconstructed curve for *P. chabaudi* (black solid line), with the corresponding inflection point (*p*_3_ = 19.09 hours) indicated by a red dotted line. Bands with around solid lines show for 95% confidence intervals for the fitted inflection point([18.68, 20.25]).

As expected from our investigation into the impact of assuming constant PMR (*R*), the best model fit returns a larger initial inoculum size (*I*_0_ = 2.53 ×10^6^) than the 10^6^ quantity used in the original study (Table S2). We compute 95% confidence intervals around the expected initial inoculum size of 10^6^ (i.e., the uncertainty due to sampling error) as [5.03 ×10^5^, 1.61× 10^6^], below the distribution of initial inoculum size estimates obtained from parametric bootstrapping (Fig. S9). This overestimation is expected given that we constrained PMRs (*R*) to be constant rather than decreasing (Fig. S7). Nonetheless, the best fitting model also recapitulates an expected feature of the early time series—that ring stage percentages would be expected to begin near 100%—despite iRBC abundance being too low to sample. Sampling from the donor mice occurred at the time of day when ring percentages were expected to be maximal [13], consistent with model-estimated ring percentages at time zero, and an initial median age of 0.71 hours (Fig. S10a). Thus, our results give no indication that model parameters or assumptions would need to be altered to describe infection dynamics prior to data collection, except that longer pre-peak time series might require a model allowing for declining PMRs.

### 3.3 Parasites exhibit shortened IDCs when misaligned to host rhythms

We first fit the 5-parameter model following the null assumption that the other treatment groups were not significantly different from the control (WT matched) group in terms of the inflection point (*p*_3_) and ring duration (*r*_*L*_). Model selection indicated that cycle length *c* was the only additional parameter required to fit dynamics in misaligned treatment groups, and the model performance was significantly better than the 5-parameter model (Fig. 5a-f, Table S3, S4, S5). In contrast to the control (WT matched) data, when parasites were misaligned to host rhythms, they exhibited shorter cycle lengths (Fig. 5g), consistent with findings reported in the original study [1]. The WT mismatched group, in which parasites experienced a 12-hour phase mismatch with the host photoperiod, had a fitted cycle length of *c* = 22.08 hours (95% CI: [22.01, 22.19]) compared to *c* = 24.00 hours for WT matched. The *Per1/2*-null TRF group, in which hosts lacked functional *Per1/2* clock genes but retained rhythmic feeding with a 12 hour phase shift from the donor mice, had a fitted cycle length of *c* = 22.51 hours (95% CI: [22.26, 22.59]). In the *Per1/2*-null all-day fed group, in which hosts lacked a functional canonical circadian clock and were fed continuously, parasites exhibited a fitted cycle length of *c* = 23.19 hours (95% CI: [22.74, 23.31]). The WT mismatched group exhibited the shortest cycle length, followed by behaviorally arrhythmic mice with time restricted feeding, suggesting that misaligned host feeding rhythms require more extreme rescheduling of IDC rhythms on the part of parasites.

**Figure 5:**
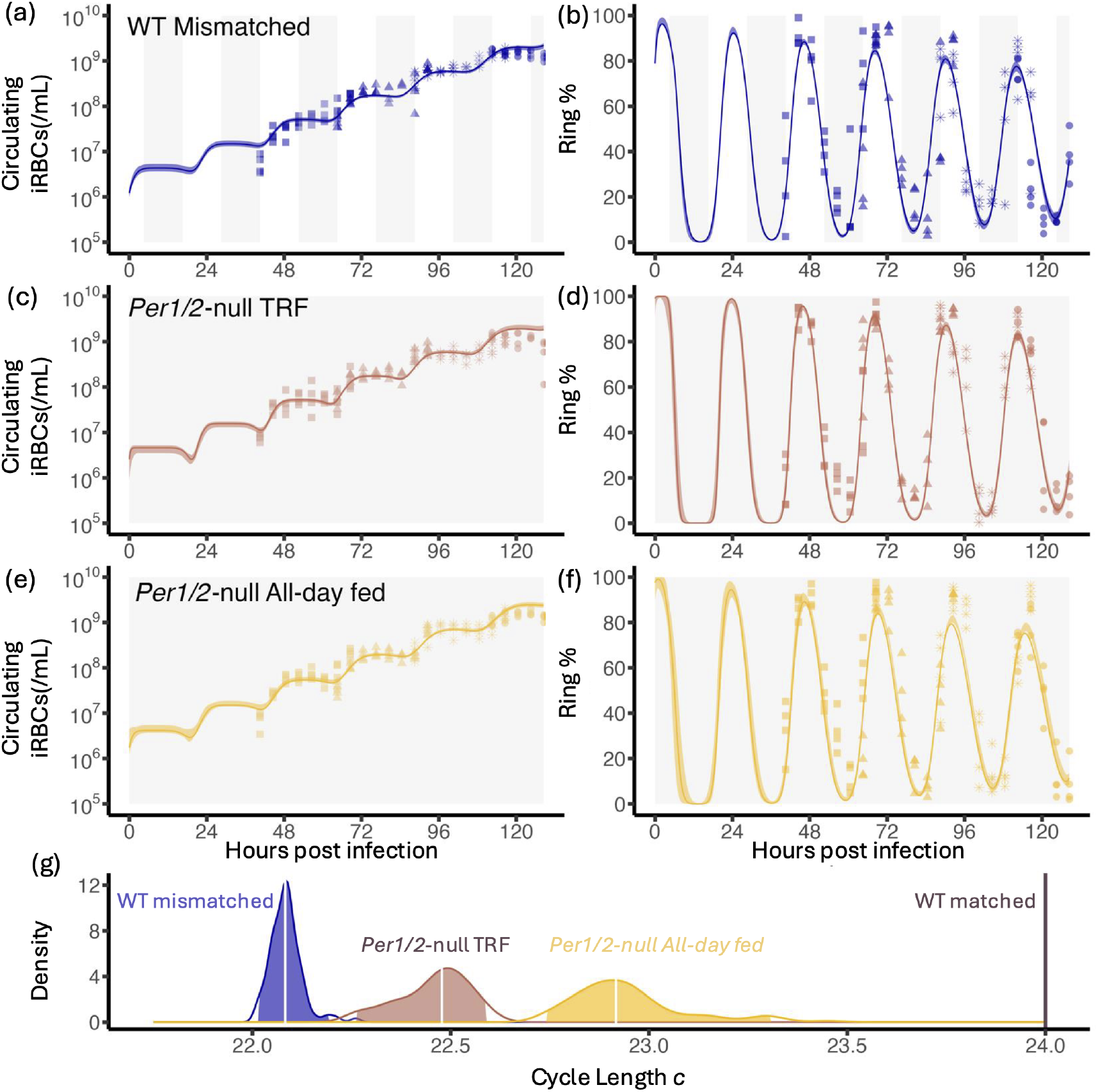
Model fits indicate that malaria parasites shorten their IDC when host rhythms are perturbed. Best-fitting model trajectories for circulating iRBC abundance and ring stage percentage are shown for the (*a-b*) WT mismatched, (*c-d*)*Per1/2*-null TRF, and (*e-f*) *Per1/2*-null all-day fed treatment groups (thick solid lines). Empirical data are shown in the background, with different point shapes indicate different mouse cohorts sampled at each time point (see O’Donnell et al. for sampling details) [1]. Grey vertical boxes indicate periods corresponding to the dark phase of the photoperiod for each treatment group. (*g*) Density distributions of fitted cycle length *c* values for each treatment group estimated via parametric bootstrapping, with white vertical lines indicating the median of each distribution. Colored regions denote the 95% confidence intervals. The WT matched cycle length (*c* = 24) is indicated by a vertical line.

Fitting either the inflection point *p*_3_ or the ring duration *r*_*L*_ gave no significant improvement in the fit to data. Accordingly, neither inflection point of sequestration curve *p*_3_ nor ring stage duration *r*_*L*_ differed significantly from WT matched values in 6 parameter models (Fig. 6a, b). These results suggest that parasites shorten their overall developmental cycle without altering early ring stage duration or the timing of sequestration.

**Figure 6:**
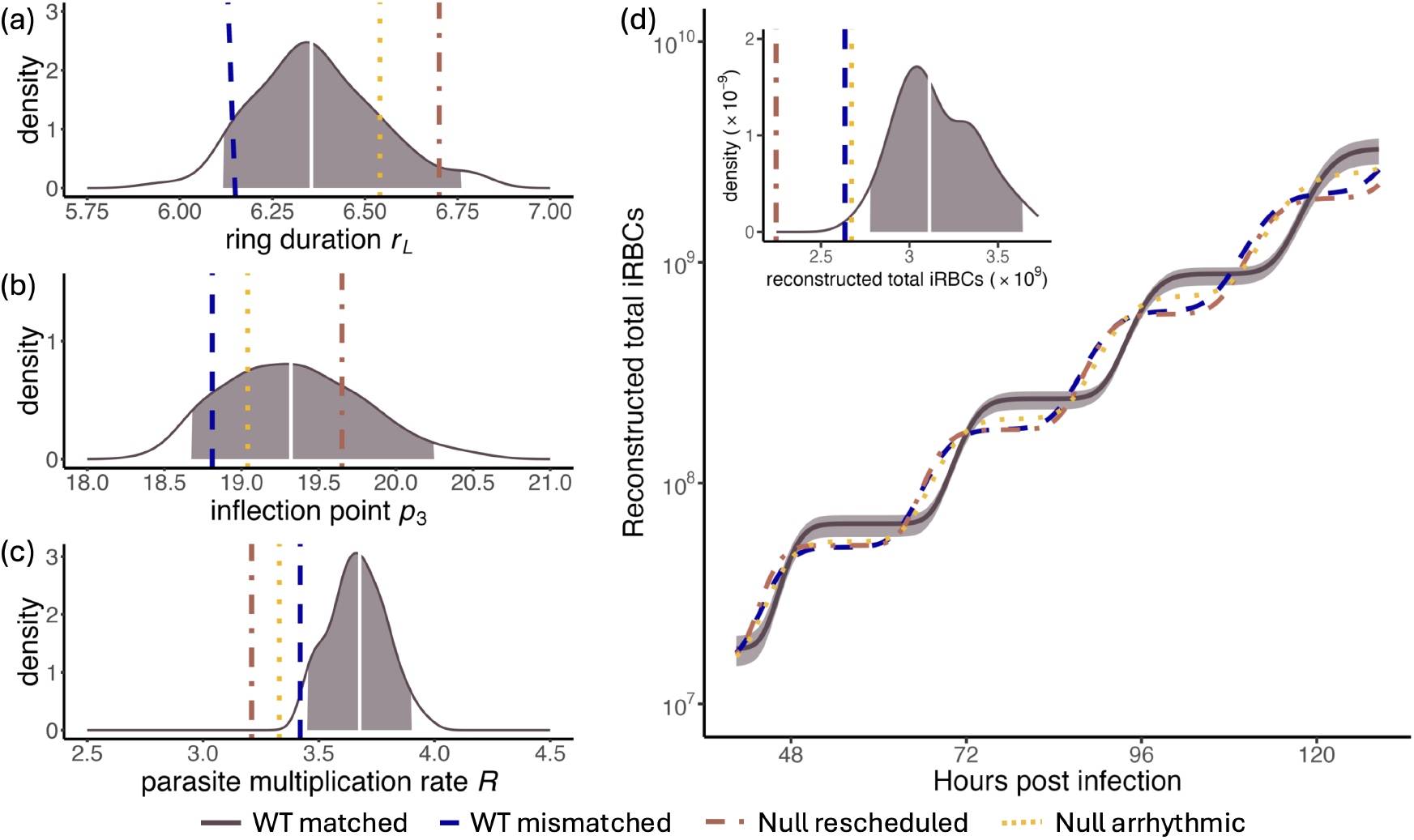
Parasites reschedule without altering ring stage duration or sequestration timing, but at the cost of reduced multiplication rates and total iRBC abundance. Density of different parameter estimates from the 7 parameter model fit to 100 synthetic time series for the WT matched data (see Methods for details), with colored area representing fitted values that fall within 95% confidence intervals for (*a*) ring duration *r*_*L*_, (*b*) the inflection point of the sequestration probability curve *p*_3_, and (*c*) PMRs *R*. Vertical lines indicate corresponding fitted values for WT mismatched (long dash), *Per1/2*-null TRF (broken line), and *Per1/2*-null all-day fed (dotted). Vertical lines indicate corresponding fitted values for WT mismatched (long dash), *Per1/2*-null TRF (dot dash), and *Per1/2*-null all-day fed (dotted). The 95% confidence intervals for each parameter are *r*_*L*_ = [6.12, 6.76], *p*_3_ = [18.68, 20.25], and *R* = [3.45, 3.90]). (*d*) WT matched infections (solid line, with band showing 95% CI from parametric bootstrapping) exhibit greater reconstructed total iRBC abundance (circulating and sequestered iRBCs) than perturbed treatment groups. The inset panel at top left shows the density distribution of the total reconstructed iRBCs at the final time point for the WT matched group, with the colored area representing the 95% confidence interval. The three vertical lines, drawn with the line types corresponding panel a-c, indicate the total reconstructed iRBCs values for each treatment group.

Similar to the WT matched results, we find that the estimated initial inoculum size (*I*_0_) significantly exceeds the expected value of 10^6^ for the perturbed treatment groups, with the *Per1/2*-null TRF treatment estimate falling significantly above the 95% confidence intervals for the WT matched group (Fig. S9). We again interpret these overestimated initial inoculum values as evidence that PMRs were decaying—especially for the *Per1/2*-null TRF treatment—rather than remaining constant as assumed in our fitted model. Both the *Per1/2*-null TRF group and *Per1/2*-null all-day fed group exhibit initial iRBC age distributions in line with expectation and with estimates for the WT matched treatment (i.e., nearly 100% early ring-stage iRBCs with median age of 0.72 hours and 1.92 hours respectively, Fig. S10c,d). Thus we find no evidence that different model assumptions would be needed to fit the early dynamics prior to data collection. In contrast, the WT mismatched group exhibits an initial median age of iRBCs of 23.04 hours, outside the bounds expected given the experimental design Fig. S10b). That deviation from expectation is difficult to explain solely in terms of declining PMRs, suggesting that different model assumptions would be needed to fit dynamics prior to the start of data collection. One possibility is an abrupt early shift in IDC duration, which could not be captured with our fitted model. Such a shift could be caused by parasites being better able to sense and respond to misaligned host rhythms in hosts with an intact canonical circadian clock. Alternately, parasites may experience a more extreme mismatch when misaligned to host feeding and other TTFL-driven rhythms, resulting in abrupt parasite mortality and changes in PMR. Data from the first days of infection would be needed to distinguish between these hypothesis, presently challenging to obtain due to the detection limits of microscopy.

### 3.4 Rescheduling impairs parasite population growth

To measure costs of rescheduling, we used the best fitting models to compare PMRs (*R* values) and reconstruct total iRBC abundance for all four treatment groups. The fitted values and confidence intervals constructed via parametric bootstrapping indicate that all perturbed treatment groups have significantly lower PMRs (*R*) than the WT matched group, meaning fewer new iRBCs produced on average per bursting iRBC (Fig. 6c). The WT mismatched group has a PMR of 3.42 new iRBCs/bursting iRBC, the *Per1/2*-null TRF group has a PMR of 3.21, and the *Per1/2*-null all-day fed group has a PMR of 3.33, all three treatment groups fall outside of the 95% confidence interval for the WT matched group where CI = [3.45, 3.90]. These estimates represent PMR per life cycle (*PMR*_*LC*_, to borrow terminology from *P. falciparum*, [35]) and therefore explicitly account for the shortened IDC observed in the treatment groups. Thus, misaligned parasites with abbreviated IDCs produce fewer new iRBCs/bursting iRBC than aligned parasites that take the full 24 hour.

In the original study [1], analysis of circulating iRBCs revealed no discernible fitness cost using generalized linear models, though the observation that parasites could not achieve greater circulating iRBC abundance by completing more IDC than in the WT matched infections over the monitoring window suggests a cost or constraint. We recreated this analysis using only pre-peak data for our revised cut-off of 128.5 hours using average circulating iRBCs to enable comparison with model-reconstructed total iRBC abundance (i.e., the sum of circulating and sequestered iRBCs, for which only one value is available per time point). We find the same results as in the original study, that circulating iRBC densities are best explained by hours post-infection alone with no need to include treatment (Table S6). In contrast, model–reconstructed total iRBC densities are best explained by the interaction of hours PI and treatment (Table S7). In all treatment groups, total iRBC abundance follows a stair-step pattern, with a discrete period of increase due to synchronized bursting followed by a plateau in numbers as parasites develop within the iRBCs they have invaded. When parasites are misaligned to host rhythms, they begin increasing earlier than the WT matched infections due to shortened IDC, but in each cycle, total iRBC numbers plateau below what WT matched parasites can achieve with a longer cycle length (Fig. 6d). Model-reconstructed estimates of total iRBC abundance show significantly reduced abundance in perturbed treatments compared to WT matched (Fig. 6d) at the peak of infection (i.e., the final timepoint we include in our analysis). Together, these results demonstrate that IDC rescheduling carries a fitness cost in the form of reduced multiplication rates and subsequently reduced total iRBC abundance within the host.

## 4 Discussion

Assessing the functional consequences of circadian rhythms is complicated by developmental sampling bias, the reality that organisms are rarely equally accessible to sampling throughout their development [16]. The relative ease of sampling the blood-borne life cycle of malaria parasites has facilitated the use of *Plasmodium* as a model system for answering fundamental evolutionary questions [36], gaining insight into the role of immunity [37], and interrogating the adaptive significance of circadian rhythms in pathogenic organisms [3]. Yet it remains challenging to obtain a complete picture of within-host infection dynamics due to other aspects of malaria biology, since parasites essentially disappear behind a curtain as they mature and sequester ([14], reviewed in [20]). To peek behind the curtain, we developed model fitting approach to reconstruct within-host dynamics from observable data. Applying this approach to data from experimental *P. chabaudi* infections of mice [1], we found that parasites shorten their life-cycle length when misaligned to host rhythms, incurring a fitness cost in the form of reduced parasite multiplication rates and total iRBC abundance within the host. Notably, parasites do not alter ring stage duration or the timing of sequestration as they reschedule. By reconstructing the within-host dynamics of circulating and sequestered iRBCs, we inferred the sequestration timing of *P. chabaudi*, which closely resembles the timing of expression of sequestration proteins observed in *P. falciparum in vitro* [28]. The approach we develop here provides a way to reconstruct within-host dynamics despite sequestration and interrogate the fitness benefits of matching host rhythms.

One key challenge is that the extent of developmental sampling bias depends on the environment in which organisms find themselves. When parasites attempt to sequester within host tissues, they express specific proteins on the surface of iRBCs, which bind to corresponding host proteins [10]. The parasite component of this process can be characterized *in vitro*, particularly for species such as *P. falciparum* that can be maintained in continuous culture [31]. The host contribution has been explored in experimental rodent infections with studies using genetically modified mice (e.g., [24]). Together, these approaches have provided important insight into the molecular determinants of sequestration, but reconstructing the dynamics of sequestration *in vivo* requires modeling. Models to date have reconstructed total iRBC burden by making assumptions about IDC scheduling (e.g., from biomarker data such as PfHRP2, [26]) or the timing of sequestration [14]. We build on those approaches by developing a model-fitting approach that fits—rather than assumes—the timing of IDCs and sequestration. We show that it is possible to infer aspects of life-cycle scheduling from observable dynamics, rather than specifying them *a priori*. Since the data used here are rarely obtainable (especially from human infections), a key area for further work is to determine how sparsely-sampled time series can be and still return useful information about parasite schedules and sequestration timing.

Despite their different life cycle lengths, *P. chabaudi* and *P. falciparum* exhibit remarkably similar sequestration probability curves, suggesting at least two hypotheses. One possibility is that sequestration timing may be governed by intrinsic biological constraints rather than scaling with IDC. If parasites require a fixed amount of time and resources to synthesize and traffic sequestration ligands to the RBC surface, that could lead to conserved sequestration timing across species despite differences in IDC. If true, conservation of developmental sampling bias would greatly simplify efforts to compare biology, including developmental schedules, across *Plasmodium* species. An alternative hypothesis is that the timing of sequestration varies within species, including across *P. chabaudi* strains, and that the strain used in these experiments (DK) sequesters unusually late in development. In support of this latter possibility, DK is often used in experiments since it is known to be one of the least virulent (harmful) strains among those common used [38]. Sequestration is thought to contribute to morbidity in human infections [19, 20], with similar patterns reflected in experiments with another rodent malaria species, *P. berghei*. In infections of genetically modified *rag1*^*−/−*^ mice, *P. berghei* parasites exhibit little to no sequestration so that mature iRBCs are cleared more efficiently by innate host immunity, and induce far less morbidity compared to infections in WT mice [25]. If DK sequesters unusually late, that might facilitate more efficient immune control of infections and account for its low virulence. By providing a means to reconstruct the timing of sequestration from *in vivo* data, our model-fitting approach can be used to identify within-and across species variation in sequestration timing and implications for virulence.

Our results show that timing of different developmental stages can be estimated for individuals from time series of populations. Some caution is warranted in interpreting model-fitted ring durations at face value due to the subjectivity inherent to categorizing a continuous developmental process into discrete morphologically identifiable stages, but our estimate of approximately 6 hours for ring stage development is consistent with previous estimates of 12 hours required for parasite populations to complete this phase of development [13]. This is because synchrony is imperfect, so populations would be expected to require more time to complete a given stage of development than would individuals. Critically, our approach does not require that stage percentage data are objective, only that staging has been performed consistently over the course of an experiment. Thus, our approach represents an alternative to current methods for estimating parasite age distributions from human malaria infections, which rely on gene expression data from artificial culture [39] that may not encompass important aspects of the within-host environment [27]. Our model fitting approach represents a much-needed alternative that can be used to interrogate variation in developmental schedules despite important limitations in methods for data collection: There is no guarantee that the timing of development is the the same *in vitro* and *in vivo*, methods for continuous artificial culture are lacking in *P. chabaudi* and most malaria species, and individual iRBCs cannot be tracked through their development *in vivo*. This challenge is not unique to malaria parasites, but applies whenever information is desired about the developmental timing of organisms in their natural habitat, even if individuals are difficult to sample at certain times in their development.

We find that across all treatment groups, parasites maintain a similar ring stage duration and sequestration timing even when the overall IDC is shortened to reschedule with host rhythms. One possibility is that mature parasites could have better capacity to sense time of day from within-host cues. Another possibility is that this pattern arises due to fundamental constraints on developmental timing, since early developmental steps are required for later developmental processes to occur. Thus, it may be lower risk for parasites to maintain flexibility in later developmental timing so as to maintain synchrony with the rest of the cohort or at the right time of day. This type of scheduling would be similar to other organisms like periodical cicadas, which appear synchronize emergence by coordinating later life stages with environmental cues [40].

A key insight from our analysis is that circadian rescheduling in malaria parasites carries a fitness cost that was not apparent without model-reconstructed within-host dynamics. By reconstructing the full within-host parasite population—including sequestered stages—we show that misaligned parasites exhibit reduced parasite multiplication rates, indicating a significant fitness penalty. Thus, perturbing host rhythms—or tricking parasites into speeding up their IDC—could hinder parasite proliferation and augment the impact of immunity and drug treatment. Applying this framework also reveals that parasites reschedule their developmental cycles by shortening later stages of the IDC, that sequestration timing is robust to parasite-host misalignment, and that there are intriguing similarities in sequestration timing across the very few species for which information is available. Sequestration in malaria parasites represents an extreme example of developmental sampling bias, but the same problems will apply across taxa. Together, these findings demonstrate that modelfitting can reconstruct unobservable dynamics and identify the trade-offs that constrain the evolution of circadian rhythms in pathogenic organisms.

## Author contributions

M.A.G., Z.C., and L.A.N. co-developed fitting methods. Z.C. performed data analysis. L.A.N. developed the optimization algorithm and R package. M.A.G. and Z.C. drafted the manuscript with input from L.A.N, A.J.D. and S.E.R..

## Data availability

Data are publicly available through the original experimental study [1] at Edinburgh DataShare repository. the The supporting code is archived on GitHub repository (https://github.com/greischarlab/ParaHostEnv), and the package developed and used in this study is archived on Zenodo (https://doi.org/10.5281/zenodo.19206287).

## Acknowledgements

We thank members of the Reece and Greischar labs as well as Nicole Mideo and Lauren Childs for useful discussion. We also appreciate invaluable advice from Stephen Ellner regarding winnowing optimization.

