## supplement for "Keeping time with the host: reconstructing the developmental rhythms of malaria parasites"

### Supplemental information

#### S1 Supplemental methods

##### S1.1 Estimating cut-off time of the acute phase of infections for each treatment group

We smoothed circulating iRBC abundance using a cubic spline (*smooth.spline* function in **R**). By default, that method uses penalized least squares, and we set the degrees of freedom = 7, i.e., one less than the number of days sampled).

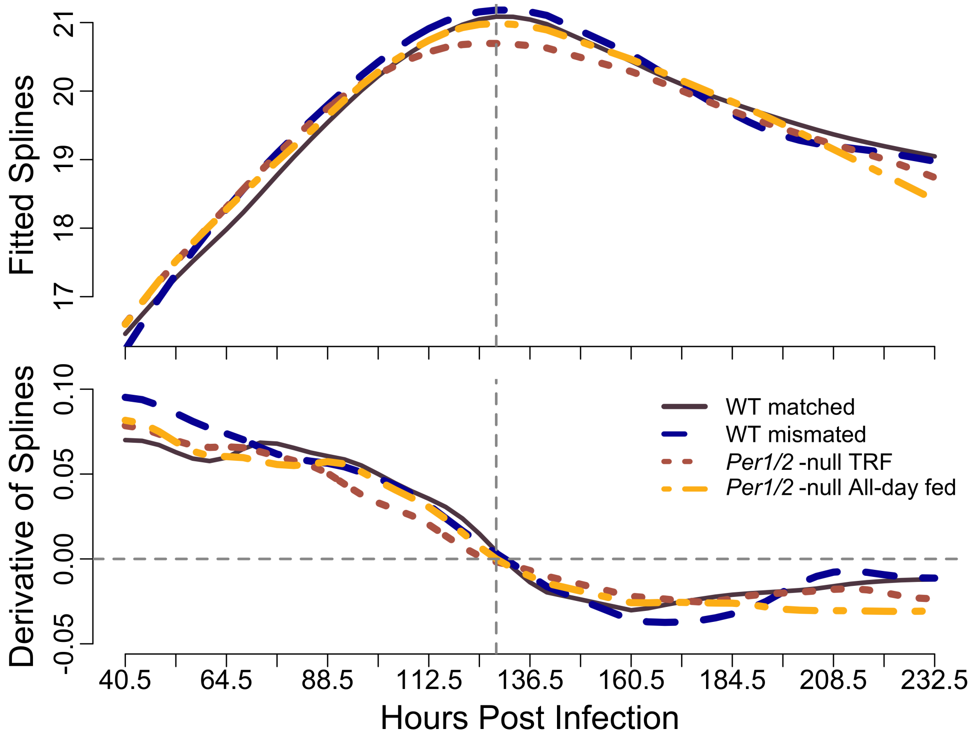


Figure S1: **The derivatives of parasite abundance for each treatment groups reach 0 at 128.5 hours post infection.** The derivatives for parasite abundance data at each sampling time point is calculated using spline (WT Matched group is represented with the black solid line, the WT mismatched group is represented with the blue dashed line, the *Per1/2*-null TRF group is represented with the brown dotted line, and the *Per1/2*-null all-day fed group is represented with the sage dot-dashed line). The horizontal grey dashed line marks for where the derivatives of fitted splines of the circulating iRBCS equal to 0, and the vertical grey dashed line marks for the corresponding time point (128.5 hours).

#### S1.2 Optimization algorithm

To locate best fit parameter values, we used a winnowing optimization method that employs multiple rounds of the Nelder–Mead algorithm implemented in the **R** package `nloptr`. We used this winnowing method to avoid poor fits due to local minima and because this method produced more robust fits to our datasets than other, simpler methods like Latin Hypercube sampling. The winnowing optimization method works by first dividing the parameter space into progressively smaller hypercubes ("boxes"), each smaller than the previous along each parameter axis by a set percentage (0.1% unless otherwise noted). The largest box encompassed the full plausible parameter space, and we reduced the parameter space by 0.1% along each parameter axis. Within each box, we randomly chose 1000 parameter sets to evaluate the objective function. The best parameter set (i.e., the set with the smallest SSE) from each box was then selected as the initial guess for coarse optimizations (maximum 1000 evaluations allowed, relative tolerance to changes $10^{-4}$). Of these we retained the best 100 (500 for the 7-parameter-model) for refined optimizations (maximum 5000 evaluations, relative tolerance $10^{-6}$). Finally, we retained the best 20 for polished optimization (maximum $10^4$ evaluations, relative tolerance $10^{-8}$). The best fit parameters were chosen as the set with the lowest SSE from the polished optimizations. Given the expanded parameter space to search for the 7 parameter model, we required the algorithm to search more boxes with greater sampling intensity. Specifically, each successive box was 0.01% rather than 0.1% smaller for the 7 parameter-model, with 5000 initial evaluations of the objective function rather than 1000. We retained the best 500 parameter sets from the refined optimizations to initiate polished optimizations.

**Implementation details.** All plots and data analyses were conducted in **R** version 4.4.2 [41]. For computational efficiency, the objective function used to calculate the sum of squared errors for each parameter combination during optimization was implemented in C++ (clang-1600.0.26.6) [42]. We implemented the winnow optimization method in the **R** package `estimatePMR` and used version 1.0.1 in our analyses [43].

#### S1.3 Parametric bootstrapping

Following the approach of [34], we used parametric bootstrapping to generate 100 synthetic datasets for the treatment groups. Because parasite density was measured from different mice at each time point, the true within-host mean–variance relationship cannot be directly observed. We therefore estimated it empirically by calculating the mean and variance of circulating iRBC densities across mice at each time point and fitting a model to characterize the distribution of the data.

The observed mean–variance relationship followed a power law of the form $\sigma^2 = a\mu^b$ (Fig. S2), indicating that the data are well described by a Tweedie distribution from the exponential dispersion family. The fitted exponent was close to two ($b = 1.884$), so we fixed $b = 2$ for simplicity, corresponding to an overdispersed Gamma distribution. Refitting the model under this assumption yielded an estimated dispersion parameter of $a = 0.106$.

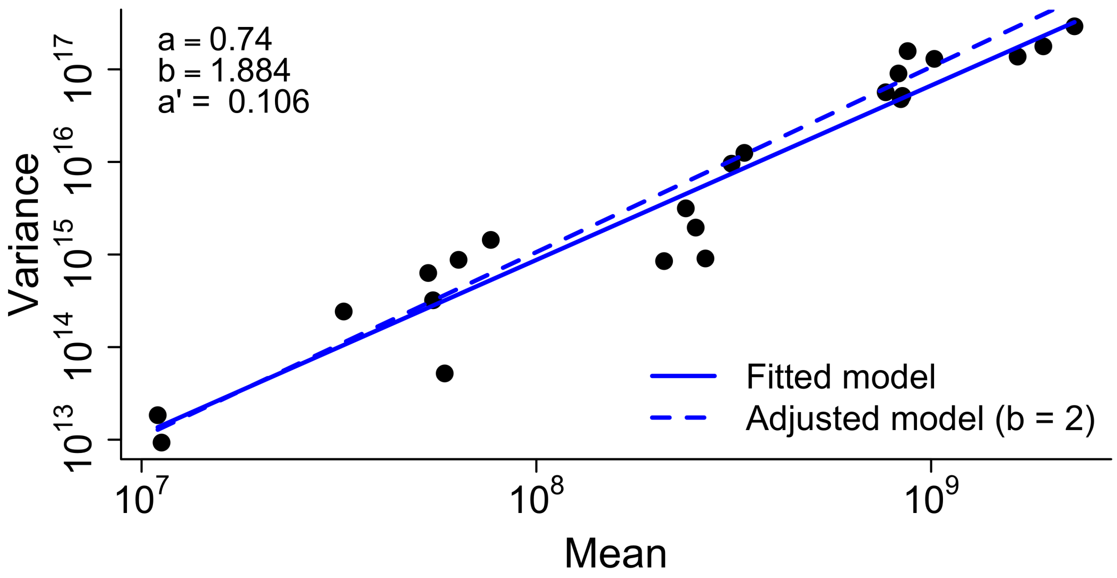


Figure S2: **The variance to mean relationship for parasite density follows a power relationship.** The black points are the means and variances calculated for data collected from each time point of the WT matched dataset. Solid blue line is the best fit assuming a power relationship between the mean and variance ($a = 0.74$, $b = 1.884$). The dashed blue line is the fit assuming the slope of the power relationship b = 2. Parameter estimate for the dispersion parameter is $a' = 0.106$ assuming a fixed slope.

Using the estimated mean–variance relationship, we generated 100 synthetic datasets for each treatment group, effectively simulating 100 independent realizations of the same experiment. These replicated datasets allowed us to repeatedly fit the ODE model and thereby estimate the sampling distributions and confidence intervals of the fitted parameters.

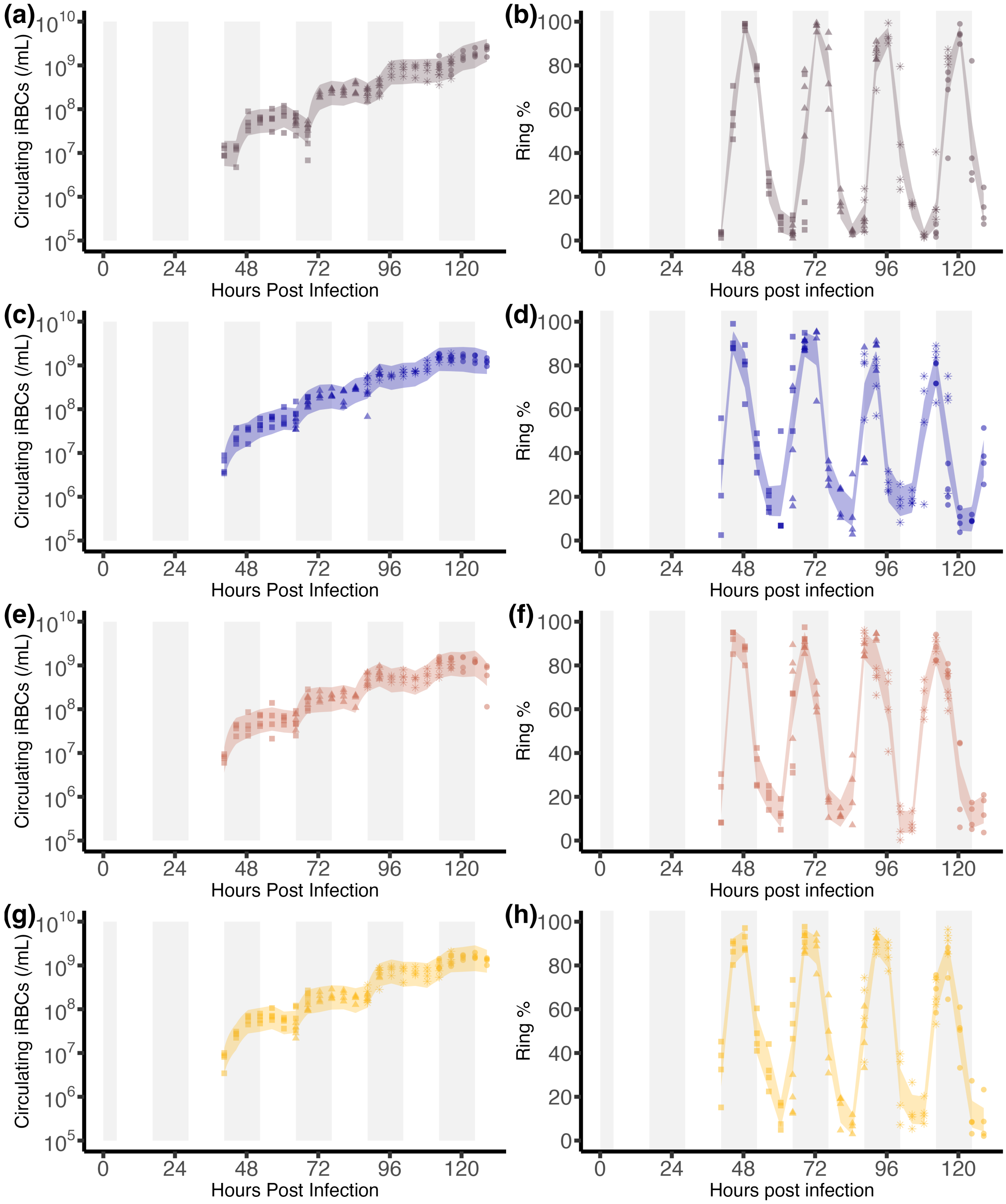


Figure S3: **Confidence intervals of the simulated iRBCs and ring percentage data represents the distribution of the parametric bootstrapped datasets.** Dots show the original dataset while different point style represents different mice cohort that the sample has been collected from. Circulating iRBCs abundance data with bands at lower transparency shows the 95% confidence interval of the 100 simulated datasets of the (a) WT matched group, (c) WT mismatched group, (e) *Per1/2*-null TRP group, and (g) *Per1/2*-null all-day fed group, with corresponding ring percentage data in (b), (d), (f), and (h). Gray rectangles in the background indicate the dark phase of the light-dark cycle for the WT matched group, plotted across all treatment groups for ease of comparison.

### S2 Supplemental figures

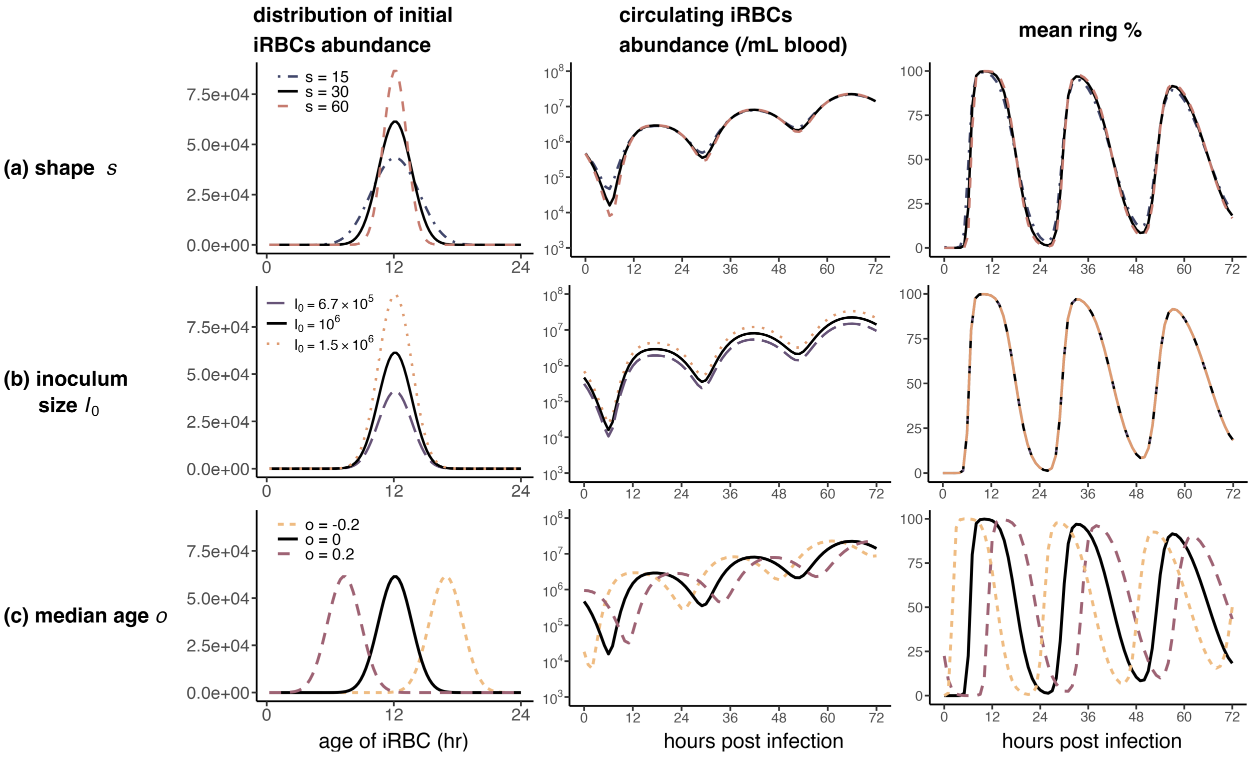


Figure S4: **Altering (a) the shape of the initial iRBC age distribution, (b) the inoculum size, or (c) the median age of the initial iRBC population has minimal impact on circulating iRBC and ring stage percentage dynamics.** Left panels show the initial iRBC age distributions, and middle and right panels show the corresponding circulating iRBC abundance and ring stage percentage dynamics, respectively. Unless otherwise stated, parameters are $p_1 = 0$, $p_2 = 1$, $p_3 = 18.57$, $p_4 = 0.4484$, $n = 100$, $R = 3$, $I_0 = 1.5 \times 10^6$, $o = 0$, $s = 30$, $c = 24$, and $r_L = 6$. The default case is shown in black solid lines. Varying the shape parameter $s$ (blue dot-dashed lines for smaller $s$ value and red dashed lines for larger $s$ value) or the inoculum size $I_0$ (purple dashed lines for larger $I_0$ and yellow dotted lines for smaller $I_0$ ) does not substantially change the dynamics. Changing the offset $o$ (yellow short dashed lines for higher median age and purple long dashed lines for lower median age) shifts the dynamics earlier or later in time, reflecting younger or older initial parasite populations, respectively.

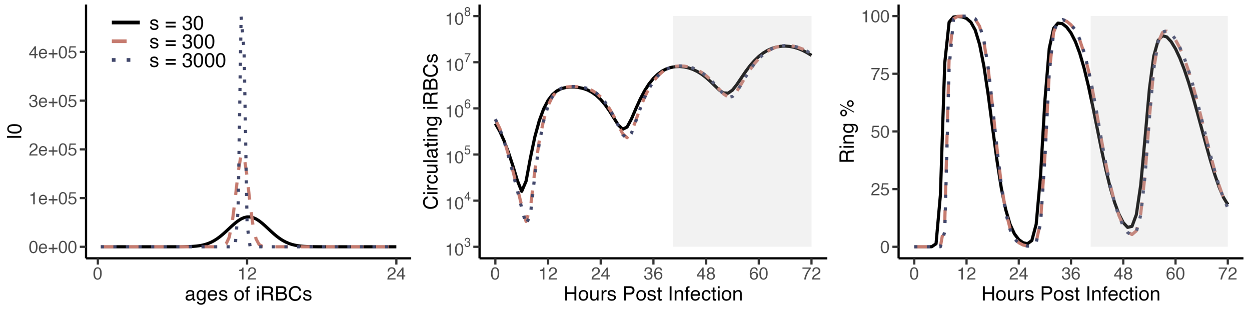


Figure S5: **Increasing shape parameter $s$ has no significant impact on dynamics of circulating iRBCs and ring percentage of the later phase of the infection.** $(a)$ The shape parameter $s$ was increased by one order of magnitude (red dashed line) and two orders of magnitude (blue dotted line) relative to the baseline value (black solid line). $(b)$ Increasing $s$ primarily alters circulating iRBCs dynamics during the early phase of infection, with enhanced sequestration during the first two parasite cycles. The predicted circulating iRBCs trajectories become nearly indistinguishable after 40.5 hours post infection where data is available (indicated by grey boxes). $(c)$ Ring-stage percentages are largely unaffected across the entire course of infection, remaining nearly identical even when $s$ is varied over multiple orders of magnitude.

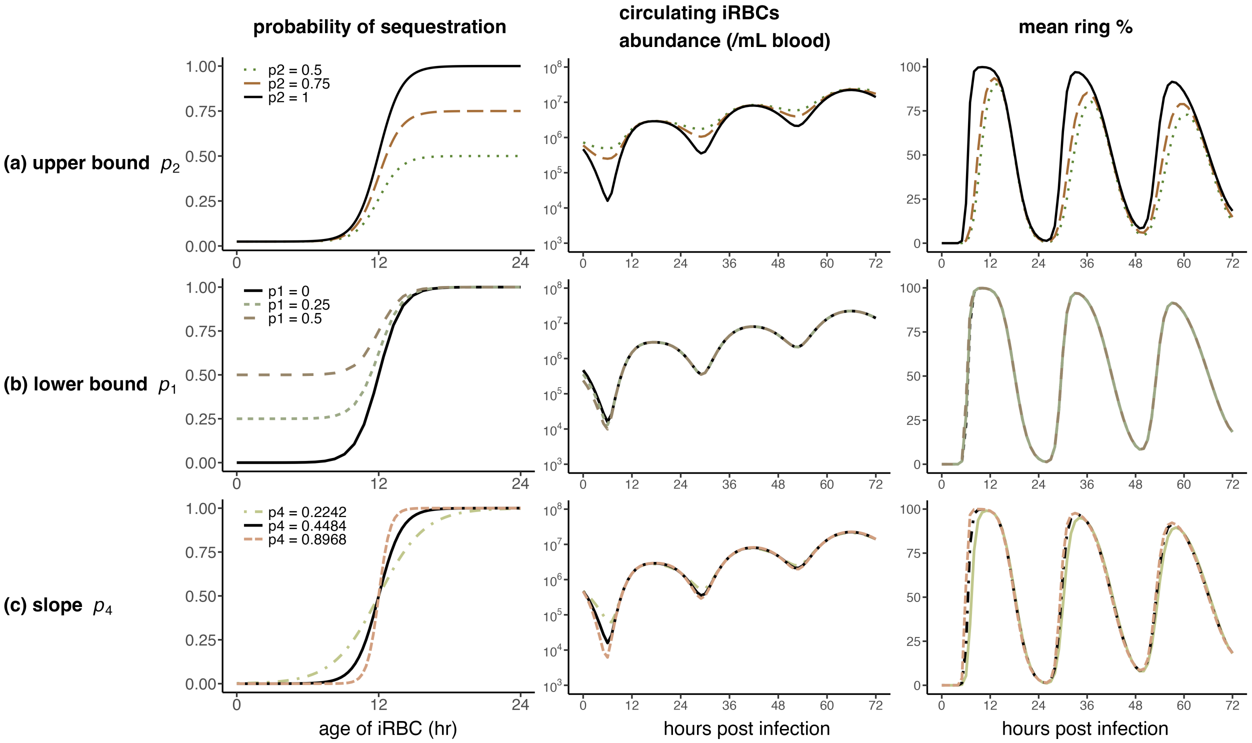


Figure S6: **Altering (*a*) the upper bound, (*b*) the lower bound, or (*c*) the slope of the sequestration probability curve has minimal impact on circulating iRBC and ring percentage dynamics.** Left panels show sequestration probability curves, and middle and right panels show the corresponding circulating iRBC abundance and ring percentage dynamics, respectively. The default case is shown in black solid lines. The upper bound $p_2$ is decreased (green dotted) or set to an intermediate value (brown dashed). The lower bound $p_1$ is increased to an intermediate (green dashed) or high value (brown dashed). The slope $p_4$ is decreased (green dot–dashed) or increased (brown dashed). Unless otherwise stated, parameters are $p_1 = 0$, $p_2 = 1$, $p_3 = 18.57$, $p_4 = 0.4484$, $n = 100$, $R = 3$, $I_0 = 1.5 \times 10^6$, $o = 0$, $s = 30$, $c = 24$, and $r_L = 6$.

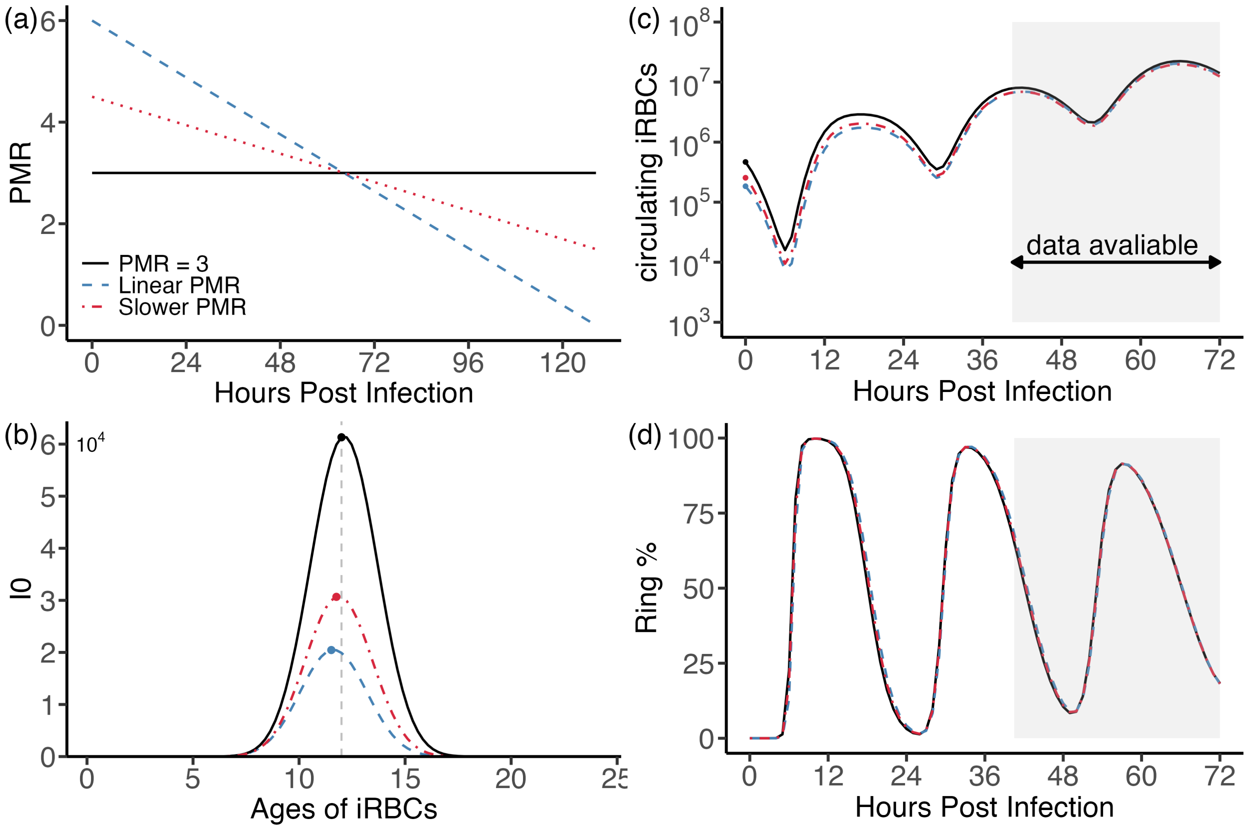


Figure S7: **A decaying PMR is associated with a higher fitted inoculum size $I_0$ while preserving observable infection dynamics.** (*a*) To fit PMR during the pre-peak phase of infection, we considered two biologically plausible modeling assumptions: (i) a constant PMR (PMR = 3, black solid line), and (ii) a time-varying, decaying PMR (steep decay: blue dashed line, slower decay: red dotdashed line), motivated by the pattern seen in Fig. S1. The decaying PMR was constrained to have the same mean value as the constant PMR over the fitted time window. We then evaluated whether these alternative assumptions lead to distinct predictions for circulating infected red blood cell (iRBC) dynamics and ring-stage proportions. (*b*) The inferred initial iRBC population size $I_0$ declines when PMR is decaying and the median age of the iRBC population is shifted earlier as well. Consequently, (*c*) the predicted circulating iRBC trajectories become nearly indistinguishable after 40.5 hours post infection where data is available (indicated by grey boxes), and (*d*) the predicted ring-stage proportion dynamics are identical under both assumptions.

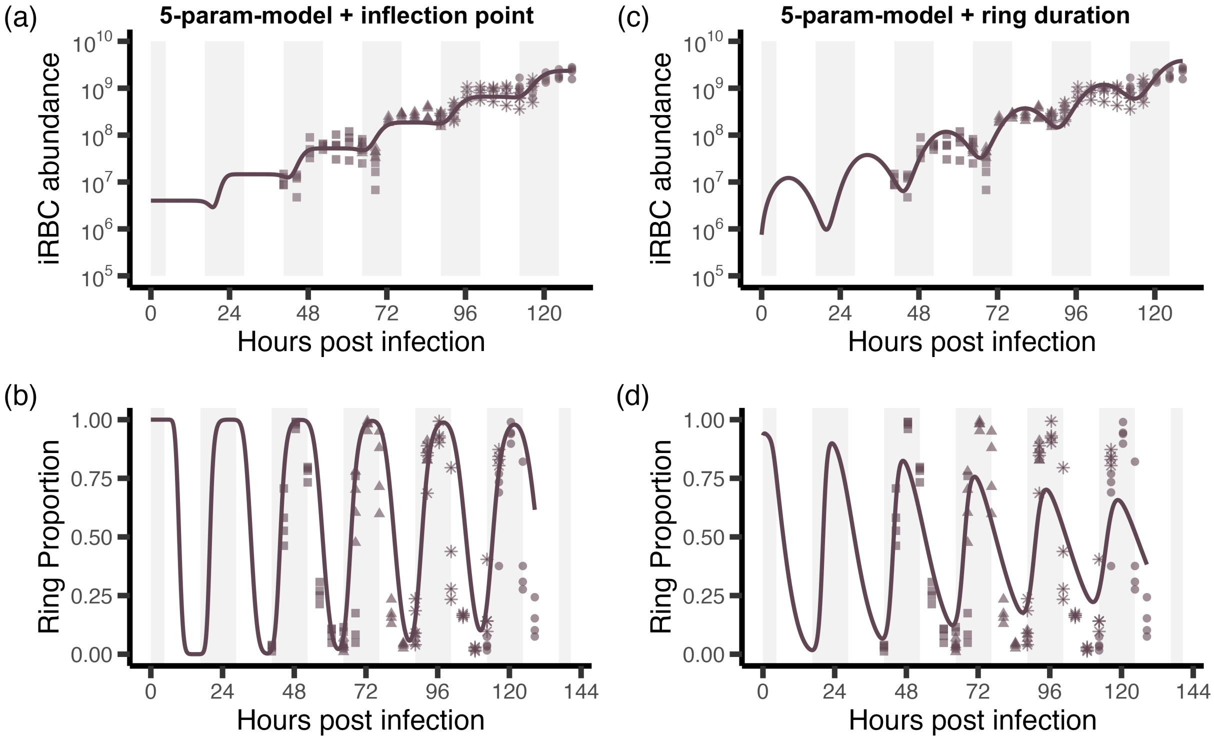


Figure S8: **Model fits to the WT Matched group indicate that inflection point $p_3$ and ring duration $r_L$ jointly improve performance, supporting their simultaneous estimation.** ($a$) Fitted circulating iRBC abundance and ($b$) ring percentage using the basic 5-parameter model augmented with the inflection point $p_3$. Solid lines denote model fits, and points represent the observed data, with different point shapes indicating distinct mouse cohorts. The grey shaded region indicates the host dark phase. ($c$) Fitted circulating iRBC abundance and ($d$) ring percentage using the basic 5-parameter model augmented with ring duration $r_L$.

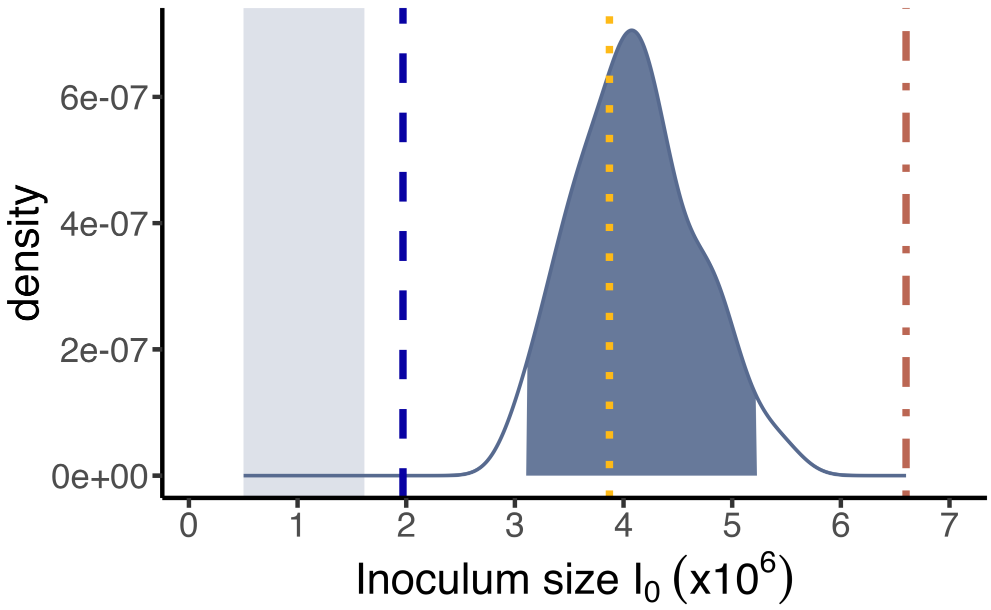


Figure S9: **Confidence intervals of fitted inoculum size $I_0$ is larger than the empirically reported value.** The blue solid line represents the density of fitted inoculum size $I_0$ of the 100 synthetic datasets, where the shaded area underneath the curve indicates the range of values that fall within the 95 percent confidence interval ([$3.11 \times 10^5$, $5.22 \times 10^6$]). Vertical lines indicate corresponding fitted values for WT mismatched (long dash), *Per1/2*-null TRF (broken line), and *Per1/2*-null all-day fed (dotted). The transparent shaded area shows the 95% confidence intervals around the expected initial inoculum size of $10^6$ (i.e., the uncertainty due to sampling error) as [$5.03 \times 10^5$, $1.61 \times 10^6$], below the distribution of initial inoculum size estimates obtained from parametric bootstrapping.

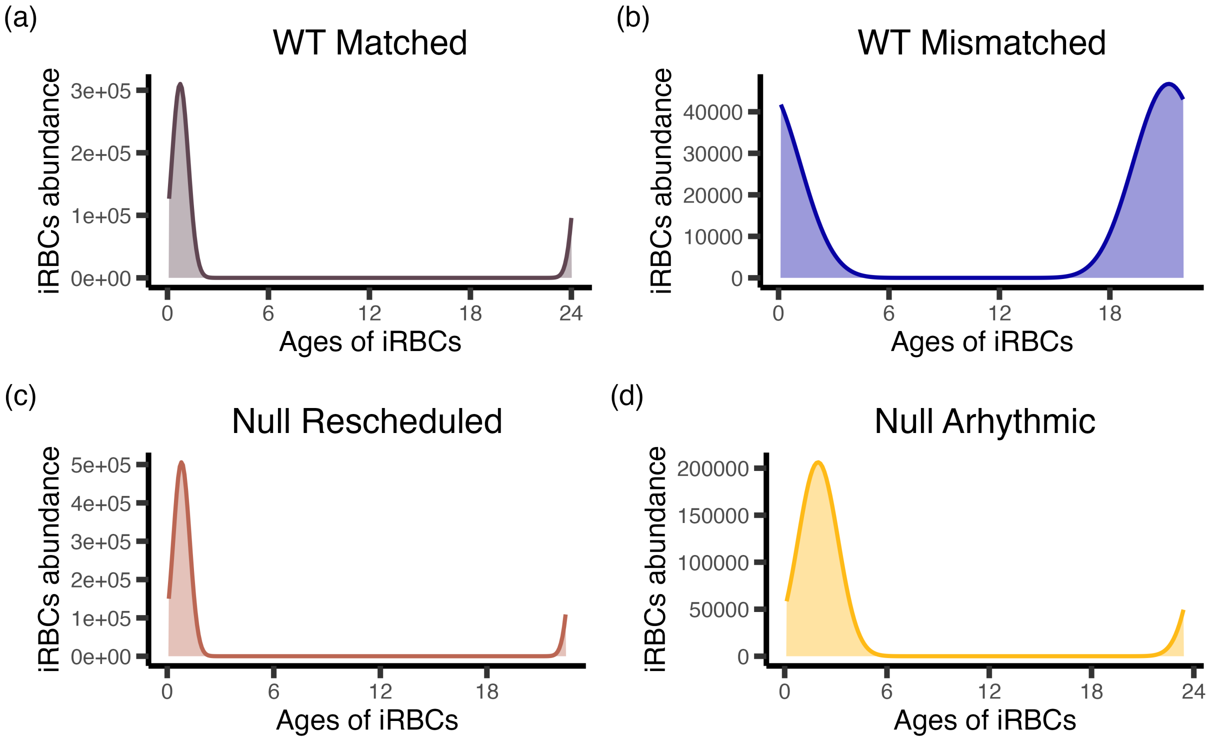


Figure S10: **Distinct initial iRBC age distributions were inferred for each treatment group under their respective best-fitting models.** ($a$) WT matched group: 5-parameter model $+r_l+p_3$. ($b$) WT mismatched group: 5-parameter model $+c$. ($c$) *Per1/2*-null TRF group: 5-parameter model $+c$. ($d$) *Per1/2*-null all-day fed group: 5-parameter model $+c$.

### S3 Supplemental tables

Table S1: Parameters used in the blood stage model of parasite development

| **Parameters** | **Fitted Range** | **Unit** |
|---|---|---|
| $R$ | $1-20$ | new iRBCs/burst/IDC |
| $c$ | $20-28$ | hr |
| $n$ | $4-500$ | - |
| $I_0$ | $\log(\min(\text{Circ iRBCs}))/10 -$ $\log(\min(\text{Circ iRBCs}))\cdot 10$ | / mL of blood |
| $o$ | $0-1$ | - |
| $s$ | $0-300$ | - |
| $r_L$ | $3-9$ | hr |
| $p_3$ | $14-22$ | hr |

Table S2: Fitted parameter values, SSE, and p-values for F test for the models tested during model selection for the WT Matched treatment group

| **Model** | $n$ | $R$ | $o$ | $s$ | $I_0$ | $c$ | $r_L$ | $p_3$ | **sse** | **p-value** |
|---|---|---|---|---|---|---|---|---|---|---|
| 5-param-model | 78 | 3.38 | 0.64 | 154.32 | $3.38 \times 10^6$ | 24* | 12* | 9.29* | 56.60 | - |
| 5-param-model + $c$ | 64 | 3.35 | 0.65 | 300 | $3.96 \times 10^6$ | 24.00 | 12* | 9.29* | 56.51 | 0.66 |
| 5-param-model + $p_3$ | 282 | 3.56 | 0.39 | 300 | $4.10 \times 10^6$ | 24* | 12* | 22.00 | 14.20 | $< 0.05$ |
| 5-param-model + $r_L$ | 125 | 3.40 | 0.63 | 7.12 | $4.75 \times 10^6$ | 24* | 3.59 | 9.29* | 10.53 | $< 0.05$ |
| 5-param-model + $p_3$ + $r_L$ | 253 | 3.68 | 0.47 | 300 | $2.53 \times 10^6$ | 24* | 6.23 | 19.09 | 6.66 | $< 0.05$ |

* Fixed default values of a parameter (not fitted). The default value of cycle length $c$ is set to 24 hours because the developmental cycle length for *P. chabaudi* is known to be 24 hours. The ring length $r_L$ is set to 12 hours as previous studies show that the population-level ring stage duration is approximately 12 hours through empirical observations. The inflection point of the probability of sequestration curve $p_3$ is set to 9.29 based on the fitted inflection point for the sequestration protein (PfEMP1) in *P. falciparum*, which is 18.5802. Since *P. chabaudi* has a life span that's half of *P. falciparum*, it is assumed to sequester twice as fast, leading to $p_3 = 18.5802/2 = 9.29$.

Table S3: Fitted parameter values, SSE, and p-values for F test for the models tested during model selection for the WT mismatched treatment group

| **Model** | $n$ | $R$ | $o$ | $s$ | $I_0$ | $c$ | $r_L$ | $p_3$ | **sse** | **p-value** |
|---|---|---|---|---|---|---|---|---|---|---|
| 5-param-model | 79 | 3.84 | 0.35 | 300 | $3.54 \times 10^6$ | 24* | 6.23* | 19.09* | 10.23 | - |
| 5-param-model + $c$ | 197 | 3.42 | 0.50 | 12 | $1.97 \times 10^6$ | 22.09 | 6.23* | 19.09* | 7.45 | $< 0.05$ |
| 5-param-model + $p_3$ | 75 | 3.82 | 0.36 | 300 | $3.68 \times 10^6$ | 24* | 6.23* | 18.81 | 10.22 | 0.73 |
| 5-param-model + $r_L$ | 79 | 3.84 | 0.35 | 300 | $3.54 \times 10^6$ | 24* | 6.15 | 19.09* | 10.23 | 0.99 |

Table S4: Fitted parameter values, SSE, and p-values for F test for the models tested during model selection for the *Per1/2*-null TRF treatment group

| **Model** | $n$ | $R$ | $o$ | $s$ | $I_0$ | $c$ | $r_L$ | $p_3$ | **sse** | **p-value** |
|---|---|---|---|---|---|---|---|---|---|---|
| 5-param-model | 133 | 3.18 | 0.30 | 300 | $5.93 \times 10^6$ | 24* | 6.23* | 19.09* | 10.40 | - |
| 5-param-model + $c$ | 244 | 2.92 | 0.46 | 200 | $6.60 \times 10^6$ | 22.51 | 6.23* | 19.09* | 6.54 | $< 0.05$ |
| 5-param-model + $p_3$ | 133 | 3.21 | 0.29 | 300 | $5.61 \times 10^6$ | 24* | 6.23* | 19.65 | 10.37 | 0.56 |
| 5-param-model + $r_L$ | 120 | 3.16 | 0.29 | 300 | $5.93 \times 10^6$ | 24* | 6.70 | 19.09* | 10.22 | 0.15 |

Table S5: Fitted parameter values, SSE, and p-values for F test for the models tested during model selection for the *Per1/2*-null all-day fed treatment group

| **Model** | $n$ | $R$ | $o$ | $s$ | $I_0$ | $c$ | $r_L$ | $p_3$ | **sse** | **p-value** |
|---|---|---|---|---|---|---|---|---|---|---|
| 5-param-model | 160 | 3.43 | 0.35 | 300 | $5.42 \times 10^6$ | $24^*$ | $6.23^*$ | $19.09^*$ | 5.73 | - |
| 5-param-model + $c$ | 378 | 3.33 | 0.42 | 12 | $3.81 \times 10^6$ | 23.19 | $6.23^*$ | $19.09^*$ | 5.33 | $< 0.05$ |
| 5-param-model + $p_3$ | 145 | 3.37 | 0.37 | 300 | $6.13 \times 10^6$ | $24^*$ | $6.23^*$ | 19.04 | 5.68 | 0.08 |
| 5-param-model + $r_L$ | 161 | 3.39 | 0.35 | 300 | $5.58 \times 10^6$ | $24^*$ | 6.54 | $19.09^*$ | 5.59 | 0.09 |

Table S6: Model comparison for pre-peak window parasite density with average circulating iRBCs data. Models were fit to $\log_{10}$(average circulating parasite density) with different combinations of predictors. df = degrees of freedom, $\log(L)$ = log-likelihood, AICc = corrected AIC, $\Delta$AICc = difference in AICc from best model, weight = Akaike weight.

| **Model description** | **df** | **log($L$)** | **AICc** | **$\Delta$AICc** | **weight** |
|---|---|---|---|---|---|
| hours PI | 3 | 19.148 | -31.9 | 0.00 | 0.892 |
| treatment + hours PI | 5 | 19.185 | -27.4 | 4.51 | 0.094 |
| hours PI * treatment | 7 | 19.784 | -23.7 | 8.19 | 0.015 |
| 1 (null model) | 2 | -67.187 | 138.6 | 170.48 | $< 0.001$ |
| treatment | 4 | -67.184 | 143.0 | 174.92 | $< 0.001$ |

Table S7: Model comparison for pre-peak window parasite density with reconstructed total iRBCs data.

| **Model description** | **df** | **log($L$)** | **AICc** | **$\Delta$AICc** | **weight** |
|---|---|---|---|---|---|
| hours PI * treatment | 9 | 97.688 | -175.2 | 0.00 | 0.967 |
| hours PI | 3 | 87.212 | -168.2 | 7.03 | 0.029 |
| hours PI + treatment | 6 | 88.658 | -164.3 | 10.85 | 0.004 |
| 1 (null model) | 2 | -85.665 | 175.5 | 350.65 | 0.000 |
| treatment | 5 | -85.631 | 182.0 | 357.14 | 0.000 |
